# Vulnerability and life-history traits correlate with the load of deleterious mutations in fish

**DOI:** 10.1101/2019.12.29.890038

**Authors:** Jonathan Rolland, Jonathan Romiguier

## Abstract

Understanding why some species accumulate more deleterious substitutions than others is an important question relevant in evolutionary biology and conservation sciences. Previous studies conducted in terrestrial taxa suggest that life history traits correlate with the efficiency of purifying selection and accumulation of deleterious mutations. Using a large genome dataset of 76 species of fishes, we show that the rate of deleterious mutation accumulation (measured via dN/dS, *i.e.* non-synonymous over synonymous substitution rate) is associated to the vulnerability, the life-history strategies, and the latitude of species. Our results, focusing on a large clade of aquatic species, generalizes previous patterns found so far in few clades of terrestrial vertebrates. These results also suggest that vulnerable species accumulate more deleterious substitutions than non-threatened ones, which give insights in how life-history traits, populations sizes and genetic risk of extinction can be tightly interconnected.

## Introduction

Determining the factors explaining why some species are more vulnerable than others is urgent given the acceleration of species extinction rates in the last decades (Barnosky et al. 2011). While it is accepted that a combination of demographic, ecological (Jennings 1998; Reynolds 2001) and genetic factors (Spielman et al. 2004) contribute to species extinction, their relative importance is still unclear.

Based on metrics such as geographical range or population size, the widely used red list IUCN index relies nearly exclusively on demographic and population dynamics criteria (Strona et al. 2014). Such an approach reliably measures the conservation status of a species under ongoing threats, but is not designed to assess the intrinsic vulnerability of a species to potential future threats (Miranda et al. 2017). To fill this gap and better manage the increasing problem of overfishing, an alternative vulnerability index has been developed for fish species: the *Fishbase* vulnerability index (Cheung et al. 2005). Instead of relying on demographic criterions, this index exploits the well-known fact that population size and population growth depends on life history and ecological species features, such as body length, longevity, fecundity or sexual maturity; as populations of large and long-lived species being smaller and less resilient than small and short-lived ones (Cheung et al. 2005).

Because the effective population size and genome evolution are related, the genomic sequence of a given species keeps track of the past variations in population size (Nadachowska-Brzyska et al. 2015). Therefore, long-term population size can be tracked down by measuring species genetic diversity (Romiguier et al. 2014) or the ratio of non-synonymous mutations over synonymous mutations (dN/dS; Popadin et al. 2007; Nikolaev et al. 2007; Romiguier et al. 2013; Figuet et al. 2016; Botero-Castro et al. 2017). In this study, we propose to test whether vulnerable species accumulate more deleterious alleles than non-threatened ones. We choose to focus our study on teleostean fishes, a large taxa with various life-history traits, for which i) several genomes have been recently sequenced (Malmstrøm et al. 2017) and ii) unmatched resources in terms of vulnerability index are available due to their economic importance for fishing activities. Furthermore, studies trying to link effective population size, life history traits and genome evolution focused so far exclusively on terrestrial species (mammals: Nikolaev et al. 2007; Popadin 2007; Romiguier et al. 2012, 2013; birds: Botero-Castro et al. 2017; reptiles: Figuet et al. 2016). These studies concluded that species with a “K-strategy” (i.e. large body size, long-lived and low fecundity) are more likely to exhibit small effective population sizes on the long-term (MacArthur & Wilson 1967). By comparing the life history traits and species vulnerability to the genome-wide efficiency of purifying selection in 76 teleostean species, we test here whether commonly assumed patterns in terrestrial habitats can be generalized to aquatic environments.

## Results and Discussion

For several decades, studies in conservation sciences have aimed to identify the causes of species vulnerability to extinction. Large body size, low fecundity and low rate of niche evolution have already been identified in previous studies as correlates of high extinction risk in mammals or birds (Fritz et al. 2009; Lavergne et al. 2013). Based on a dataset of 76 species of fish, our genomic analyses showed that the rate of non-synonymous over synonymous substitutions (dN/dS) was strongly associated to vulnerability (P = 1.68 x 10^-5^, R^2^_ajust_ = 0.24, Figure 1, Table 1). This association was found when dN/dS was estimated with *mapnh* and *PAML* and when we controlled for phylogenetic correlation (Figure 1, Table 1, Supplementary Material S2). We also found a strong relationship between dN/dS and life history traits (dN/dS and body length: P = 1.65 x 10^-5^, R^2^_ajust_ = 0.25; dN/dS and longevity: P = 3.69 x 10^-2^, R^2^_ajust_ = 0.11; Table 1), which is consistent with previous studies on terrestrial clades in mammals, birds and metazoa (Nikolaev et al. 2007; Popadin 2007; Romiguier et al. 2012, 2013, 2014; Botero-Castro et al. 2017; Figuet et al. 2016). Our analyses indicate that these results are not due to substitution saturation that could bias dN/dS estimations (Supplementary Material S3). Additional test showed that the relationship between dN/dS and vulnerability remained significant even when we removed the 15% of the longest branches that are potentially more affected by saturation (P<0.05, Supplementary Material S4).

**Figure 1.**
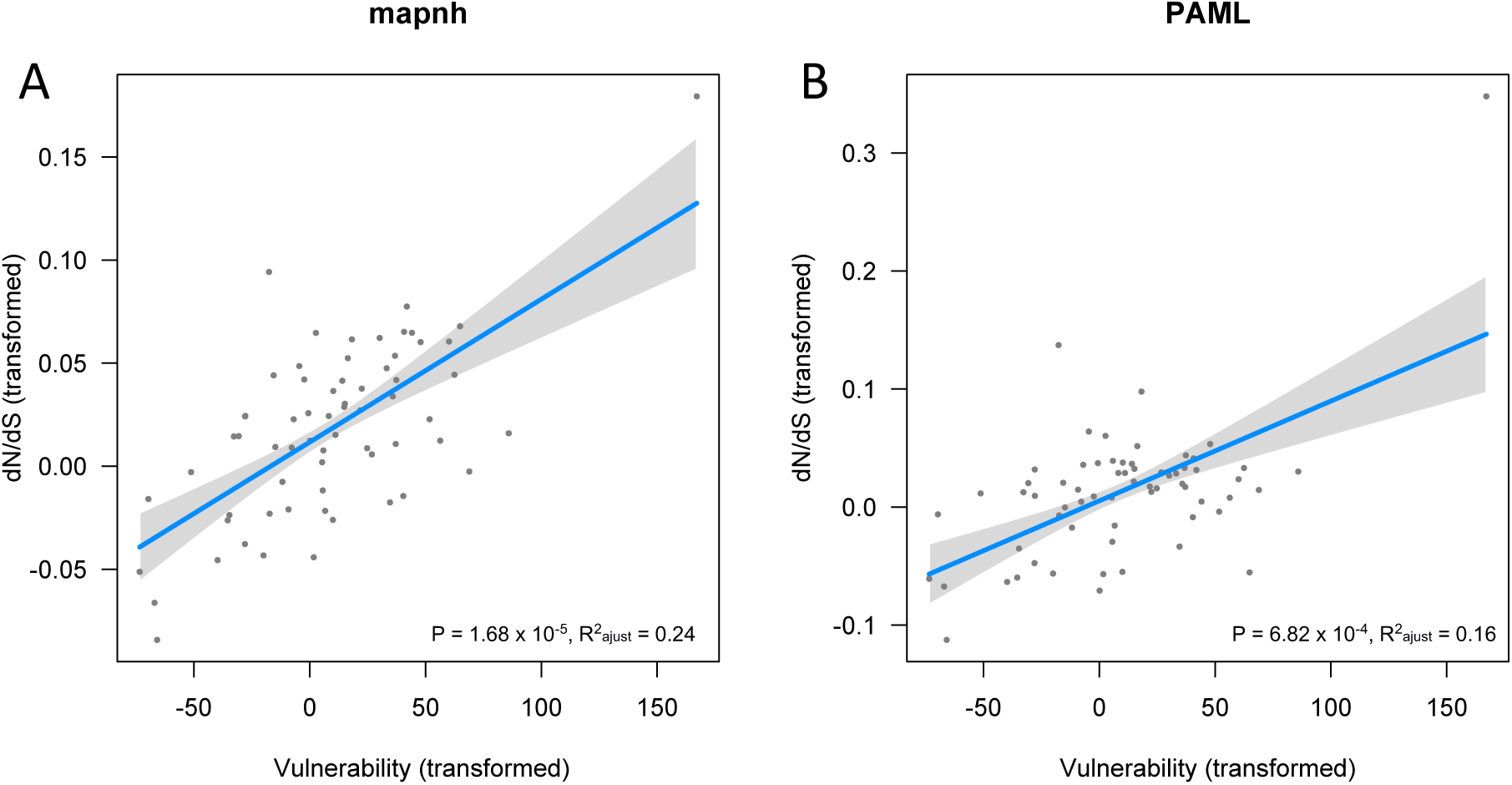
dN/dS is associated with vulnerability. dN/dS was computed using either (A) *mapnh* or (B) *PAML*. Regression line, P-values and R^2^_ajust_ of the phylogenetically generalized least squares regressions were shown on each panel respectively for all terminal branches of more than 10,000 synonymous substitutions. The shaded grey represents the 95% confidence intervals. x and y axes have been transformed in order to account for the phylogenetic relatedness between variables (script available in Supplementary material S6).

**Table 1.** Phylogenetic generalized least squares regressions between dN/dS, vulnerability, latitude, body length and longevity. Our analyses were ran with the *pgls* function which allows us to model the relationship between variables accounting for phylogenetical non-independence. dN/dS was computed using *mapnh*.

| Variable | Intercept (Std. Error) | Estimate (Std. Error) | t value | P-value | $R^2_{\text{ajust}}$ |
| --- | --- | --- | --- | --- | --- |
| Vulnerability | $5.74 \times 10^{-2}$ ($9.27 \times 10^{-3}$) | $3.72 \times 10^{-4}$ ($7.99 \times 10^{-5}$) | 4.66 | $1.68 \times 10^{-5}$ | 0.24 |
| Latitude | $5.27 \times 10^{-2}$ ($1.04 \times 10^{-2}$) | $4.21 \times 10^{-4}$ ($1.32 \times 10^{-4}$) | 3.2 | $2.17 \times 10^{-3}$ | 0.13 |
| log(Body length) | $4.51 \times 10^{-2}$ ($1.04 \times 10^{-2}$) | $7.56 \times 10^{-3}$ ($1.62 \times 10^{-3}$) | 4.67 | $1.65 \times 10^{-5}$ | 0.25 |
| log(Longevity) | $6.01 \times 10^{-2}$ ($1.20 \times 10^{-2}$) | $6.02 \times 10^{-3}$ ($2.75 \times 10^{-3}$) | 2.19 | $3.69 \times 10^{-2}$ | 0.11 |

One potential interpretation of the strong relationship found between vulnerability and dN/dS is that the accumulation of deleterious mutation is playing a role in the extinction of species. Indeed, the decrease of the efficiency of purifying selection in populations with small effective sizes (Ohta et al. 1992) has been linked to a snow-ball mechanism for species extinction known as mutational meltdown (Lynch et al. 1995). The accumulation of deleterious mutations can lead, in the long term, to a decrease of fitness resulting in a decrease of population size which ultimately will also increase the accumulation of deleterious alleles, leading to a further decrease of population size. Such negative demographic/genetic feedbacks loops toward extinction have already been proposed as a potential explanation for the local extirpation of large vertebrates (e.g. woolly mammoth, Rogers et al. 2017). In the present study, it is however difficult to disentangle causation from correlation. Indeed, the *Fishbase* vulnerability index is directly built from life-history traits, which is reflected in our data by the strong correlation between vulnerability and body size (R^2^_ajust_ = 0.60, P = 3.29 x 10^-14^). Given the fact that dN/dS correlates equally with vulnerability and other life-history traits, it is difficult to conclude that genetic factors (dN/dS) are directly causing species extinction. A more parsimonious explanation is that life-history traits (e.g. body size), vulnerability and dN/dS are all correlated simply because they are all different proxies of effective population size (Jennings and Blanchard 2004). Because they all correlate with each other, it is consequently difficult to disentangle demographic, ecological and genetic components of extinction risks, even if low population sizes, large body sizes and high rates of deleterious mutations seems to be all directly or indirectly associated to high species vulnerability. Future studies using a different metric of species vulnerability calculated independently from life history traits might help to better measure the relative contribution of ecological and genetic factor on fish extinction risk (Polishchuk et al. 2015).

Our estimate of dN/dS was also significantly higher in species distributed at higher latitude (P = 2.17 x 10^-3^, R^2^_ajust_ = 0.13, Table 1), although this relationship was not significant for the *PAML* analysis (Supplementary material S2). This result suggests that species distributed at high latitude may have generally lower or less stable population sizes. We found a significant relationship between latitude and vulnerability (PGLS, P = 5.32 x 10^-3^, R^2^_ajust_ = 0.10, Supplementary material S1) but, surprisingly, latitude was not associated with either body size (PGLS, P > 0.05, R^2^_ajust_ = 0.03) or longevity (P > 0.05, R^2^_ajust_ = −0.01). We thus hypothesize that latitude may be associated to population size by other mechanisms than an increase of body size and longevity. For example, population dynamics at high latitude might be related to other ecological factors, such as strong seasonal and long-term climatic oscillations which reduce periodically the carrying capacity of high latitude ecosystems (Dynesius and Jansson 2000). This result also implies that particular attention should be paid to vulnerable species at high latitude, given that they might have lower long-term effective population sizes, which would lead to accumulate more deleterious mutations, compared to tropical species.

The strong correlations found between vulnerability and dN/dS and between body size and dN/dS suggest that vulnerability and body size might be generally better proxies of effective population size than latitude or longevity. In theory, longevity should be also related to population size and dN/dS (Romiguier et al. 2013), but in practice the difficulty to measure longevity in the wild may lead to spurious life span measures that may blur the relationship with dN/dS. Similarly, latitude might be related to population size but species distribution is also depending on other factors such as dispersal contingencies and species interactions (Sexton et al. 2009), which are not expected to be directly related to the effective population size.

## Conclusion

Based on genomes of 76 species of fishes, our study shows that large, long-lived and vulnerable species accumulate more deleterious mutations than small, short-lived and not vulnerable species. Our results extend to aquatic environments the association between the efficiency of natural selection and species life-history strategies previously found in few clades of terrestrial vertebrates. Finally, this work also highlights for the first time a positive relationship between dN/dS and latitude, suggesting that species distributed at higher latitude might have small long-term effective population sizes, and consequently accumulates more deleterious mutations than tropical species.

## Material & Methods

### Molecular data and phylogeny

We used the alignments and the phylogenetic tree provided by the authors of Malmstrøm et al. (2017). The authors aligned 1938 exons, from 120 to 1764 bp (Supplementary Material S7), in 76 fish species using a multi-step blast procedure with 33,737 annotated zebrafish genes (from the Ensembl release 78; Cunningham et al. 2014). All alignments provided by Malmstrøm et al. are available in the zenodo dataset repository (https://zenodo.org/record/3516455#.XbjIMad7R3m) and correspond to the less strictly filtered dataset before removal of 3^rd^codon position. The phylogenetic tree was constructed with RAxML v. 8.1.17 (Stamatakis 2014) and BEAST v.2.2 (Bouckaert 2014) using 567 exons from 111 genes, with a total alignment length of 71,418 bp and 17 fossil calibrations (see Malmstrøm et al. 2017 for more details).

### Data

Body length, latitude and vulnerability data were obtained for all 76 species in *Fishbase* (http://www.fishbase.org/, Froese & Pauly 2014). Longevity was obtained for 33 species. The *Fishbase* vulnerability index is calculated from a method averaging several traits related to the rate of reproduction, such as body size and generation time (Cheung et al. 2005).

### Molecular evolution and dN/dS

We estimated the number of synonymous and non-synonymous substitutions in the terminal branches of the 76 teleost species from the alignment of 1938 concatenated genes (378,663 bp for the concatenated alignment) using the probabilistic substitution mapping implemented in the software *mapnh* (Romiguier et al. 2012; https://github.com/BioPP/testnh). The *mapnh* analysis estimates a probabilistic count of synonymous and non-synonymous substitutions. We used the ratio of the sum of non-synonymous substitutions and the sum of synonymous substitutions as our dN/dS measure. For the *PAML* analysis, we first estimated dN and dS on each individual genes (see Supplementary Material S7 for the details), and then computed the average dN and average dS by weighting with gene length.

Estimates of dN/dS can be poorly estimated when the total number of substitutions is small (short branches). We controlled for this bias by running all analyses after the removal of the terminal branches with less than 10,000 synonymous substitutions (∼15% smallest branches). We chose this threshold because it allowed to eliminate atypical short branches, which were difficult to compare with the others branches for accurate dN/dS estimations. These small branches belonged mostly to closely related species inside taxa that were over-sampled compared to the other clades, such as the *Gadinae* subfamily. We replicated the analysis with the dN/dS estimations produced by *codeml* from the *PAML* package (Yang 2007).

### Relationship between dN/dS, life history traits and vulnerability

We tested whether dN/dS was associated with life history traits and vulnerability with phylogenetic generalized least squares regressions (*pgls* function of the *R* package *caper)*. The *pgls* function allows to model a linear relationship while taking into account the phylogenetic structure of correlation between variables. For plotting the phylogenetic generalized least squares regressions, we transformed the x and y-axis in order to take into account the phylogenetic correlation in variables (code provided in Supplementary Material S6, Figure1). We then tested the robustness of the association between dN/dS and life history traits/vulnerability with two other set of analyses: one set of analyses with all species (including branches with less than 10,000 synonymous substitutions, Supplementary Material S5), and one set of analyses removing the ∼15% longest branches (branches with more than 35,000 synonymous substitution) to test whether substitution saturation was affecting our results (Supplementary Material S4). We also tested whether substitution saturation affected our results with the entropy index of substitution of Xia et al. 2003 in the software *DAMBE 7* following the procedure described in Xia & Lemey (2009; see Supplementary Material S3 for more details).

## Acknowledgments

We thank Xuhua Xia, Matthew Hahn for advices on the analyses. We also thank Dolph Schluter for providing the code to plot the phylogenetic generalized least squares regressions. JR received funding from the European Union’s Horizon 2020 research and innovation programme under the Marie Skłodowska-Curie grant agreement No. 785910 and the Banting postdoctoral fellowship (151042)

## Authors contributions

J.R. and J.R. designed the study, ran the analyses and wrote the manuscript.

---

**Supplementary Material S1.**
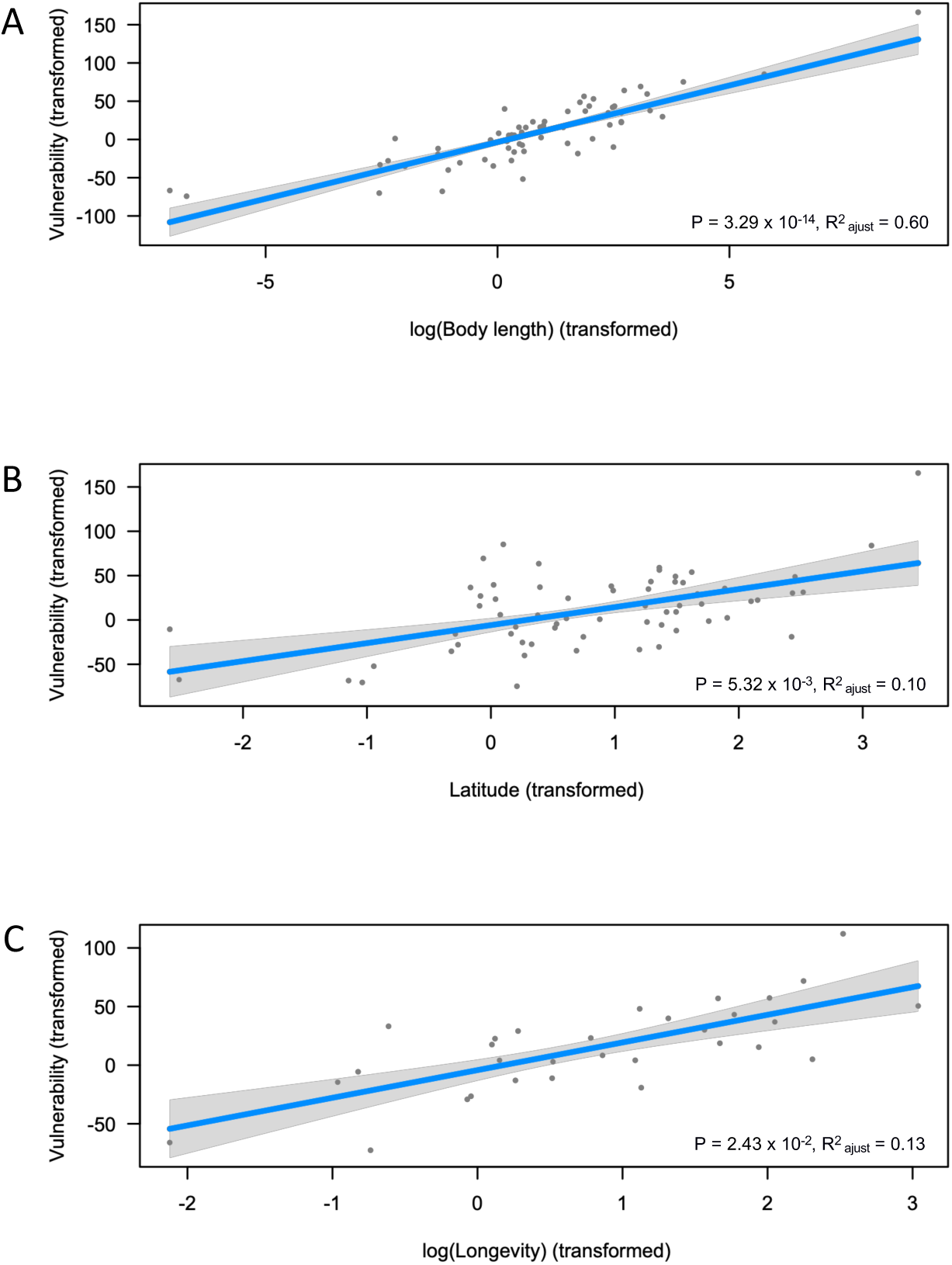
Relationship between vulnerability and (A) body length, (B) latitude and (C) longevity. Regression line, *P*-value and R^2^_ajust_ of the phylogenetic least squared regressions are shown for each variable. The shaded grey represents the 95% confidence intervals. Body length and longevity have been log-transformed, and latitude is shown in absolute values. These results were expected given that *Fishbase* vulnerability index is a metric directly computed from multiple life history traits (Cheung et al. 2005). x and y axes have been transformed in order to account for the phylogenetic relatedness between variables (script available in Supplementary material S6).

**Supplementary Material S2.**
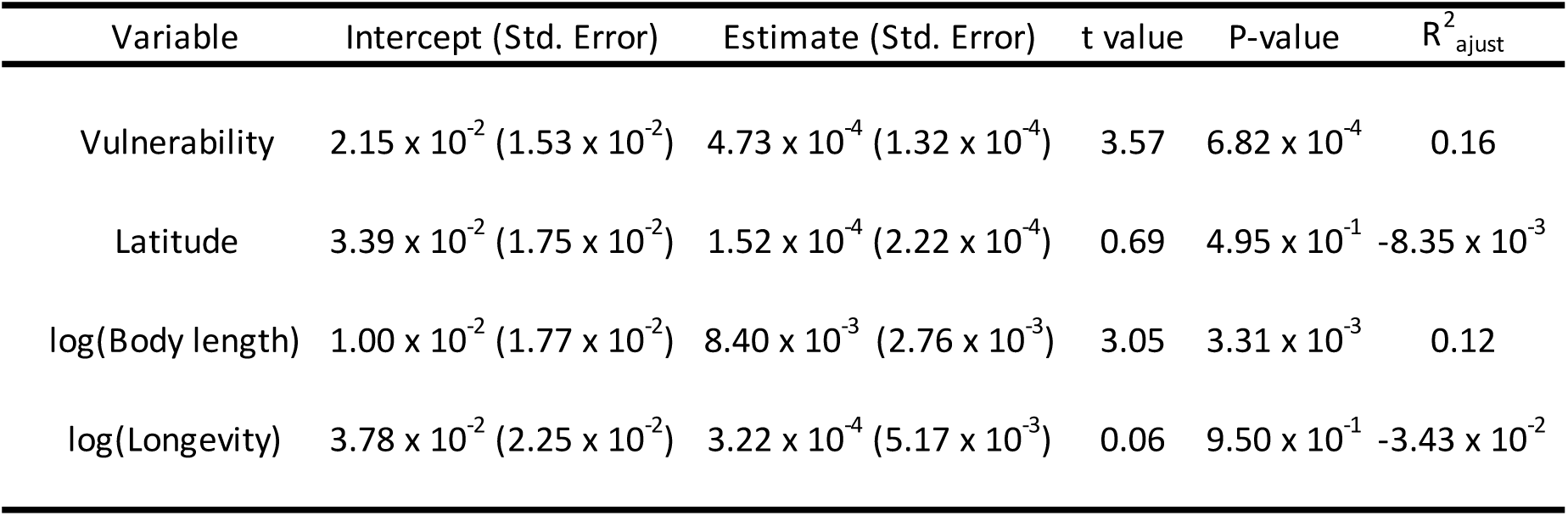
Phylogenetic generalized least squares regressions between dN/dS, vulnerability, latitude, body length and longevity. Our analyses were ran with the *pgls* function which allows us to model the relationship between variables accounting for phylogenetical non-independence. dN/dS was computed using *PAML.* We show here that our results are consistent with the results obtained with *mapnh* for vulnerability and body length.

**Supplementary Material S3.**
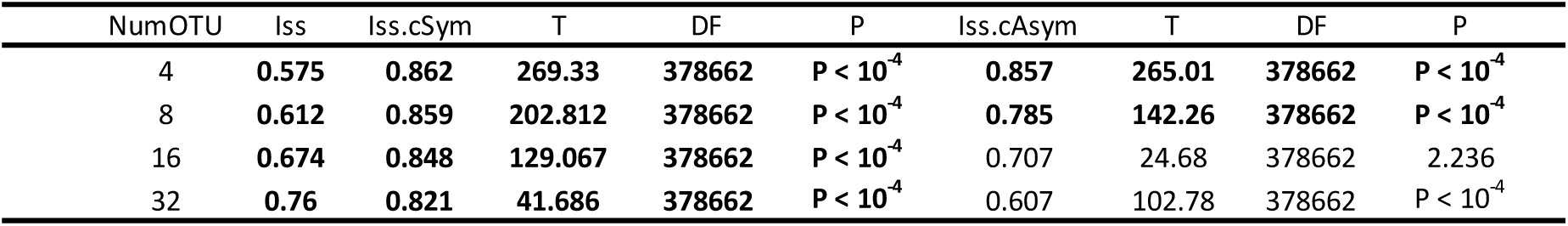
Substitution saturation analyses using the entropy index of substitution of Xia *et al.* 2003 in the software *DAMBE 7*. Estimates of the index of substitution saturation (Iss) were obtained for a proportion of invariant of P = 0.166. Substitution saturation is considered negligible only in the cases noted in bold in the table, that is to say when the Iss is significantly smaller (P<0.05) than the critical Iss value (Iss.c) at which the sequences will begin to fail to recover the true tree. Iss.cSym corresponds to the critical Iss value for symmetrical tree while the Iss.cAsym corresponds to critical Iss value for unrealistic asymmetrical tree. Because the Iss.c is based on simulations limited to 32 species in DAMBE, we have randomly sampled 200 times subsets of 4, 8, 16 and 32 species and performed the test for each subset (see NumOTU in the table). Our results suggest that substitution saturation remains very limited when the tree is symmetrical, but might be potentially problematic for recovering the true topology when the tree is extremely asymmetrical. Although our data is very unlikely falling into this category of extreme asymmetrical topology as it is considered unrealistic (Xia & Lemey 2009), we also tested whether removing long branches - with potentially more saturation - affect our results in Supplementary Material S4.

**Supplementary Material S4.**
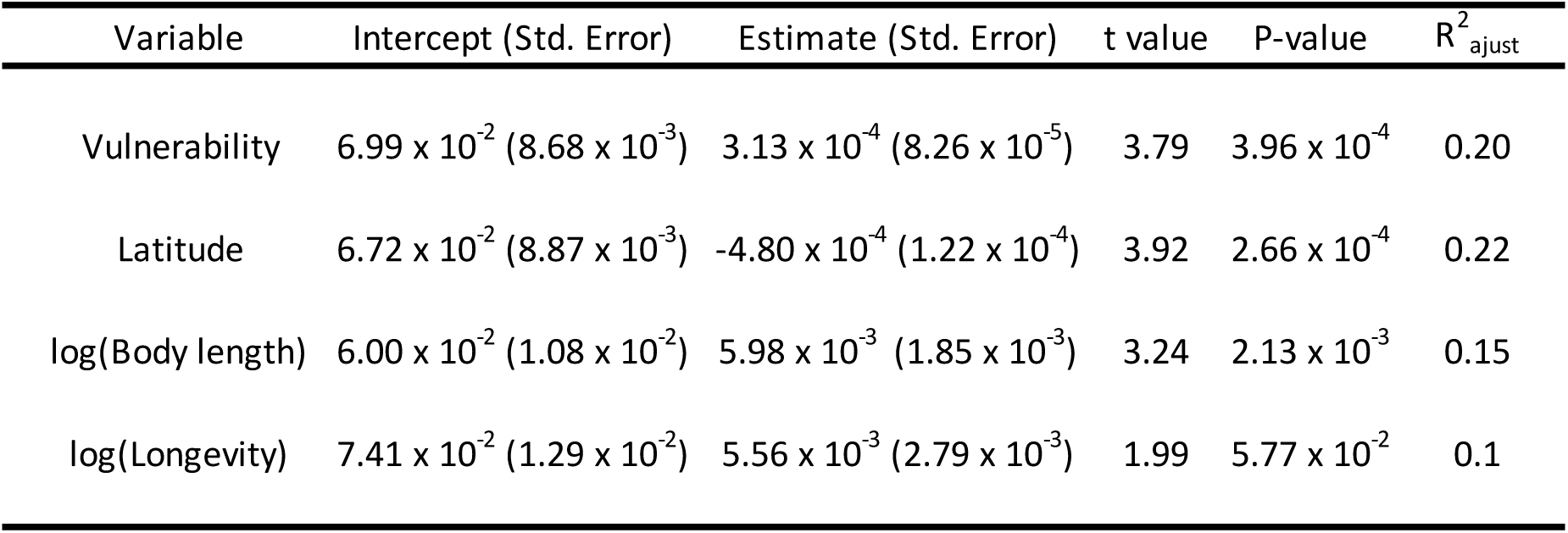
Phylogenetic generalized least squares regressions between dN/dS, vulnerability, latitude, body length and longevity obtained when long branches were removed (potentially more affected by substitution saturation). dN/dS were obtained with *mapnh*. We show here that the results presented in the main manuscript still hold when we removed the branches with more than 35,000 synonymous substitutions (∼15% longest branches; as in all other analyses, this analysis also excludes branches with less than 10,000 synonymous substitutions).

**Supplementary Material S5.**
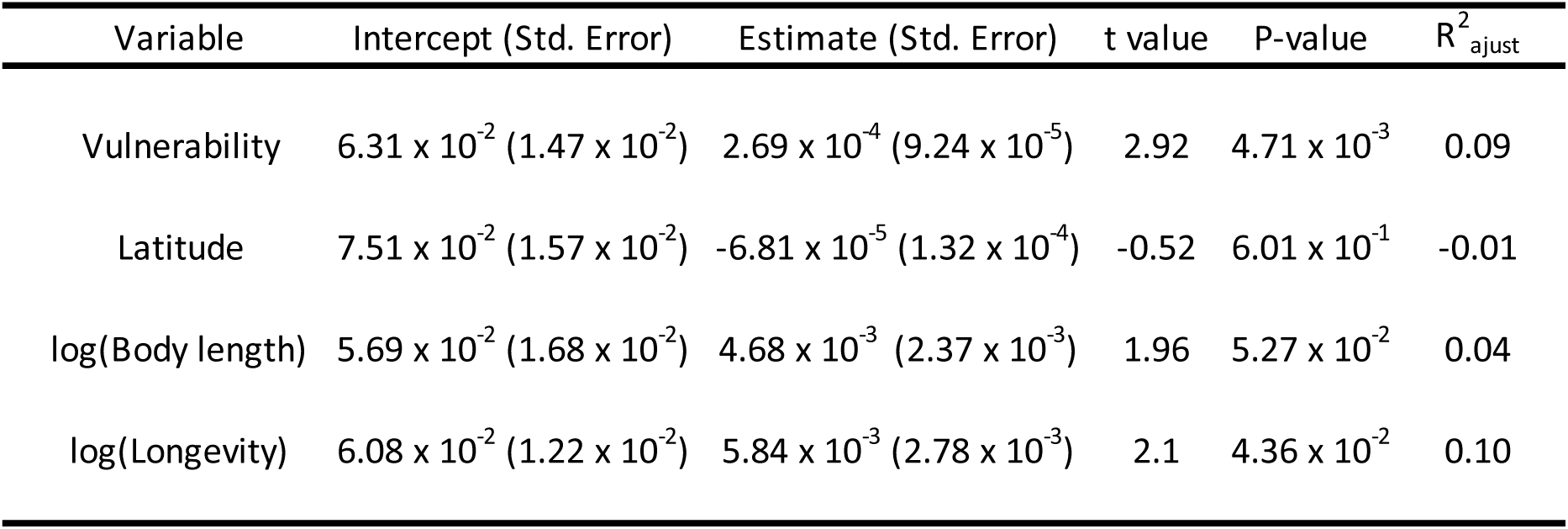
Phylogenetic generalized least squares regressions between dN/dS, vulnerability, latitude, body length and longevity obtained with all species (also include branches with less than 10,000 synonymous substitutions). dN/dS were obtained with *mapnh*. We show here that the results presented in the main manuscript still hold for vulnerability, body length and longevity.

**Supplementary Material S6.**
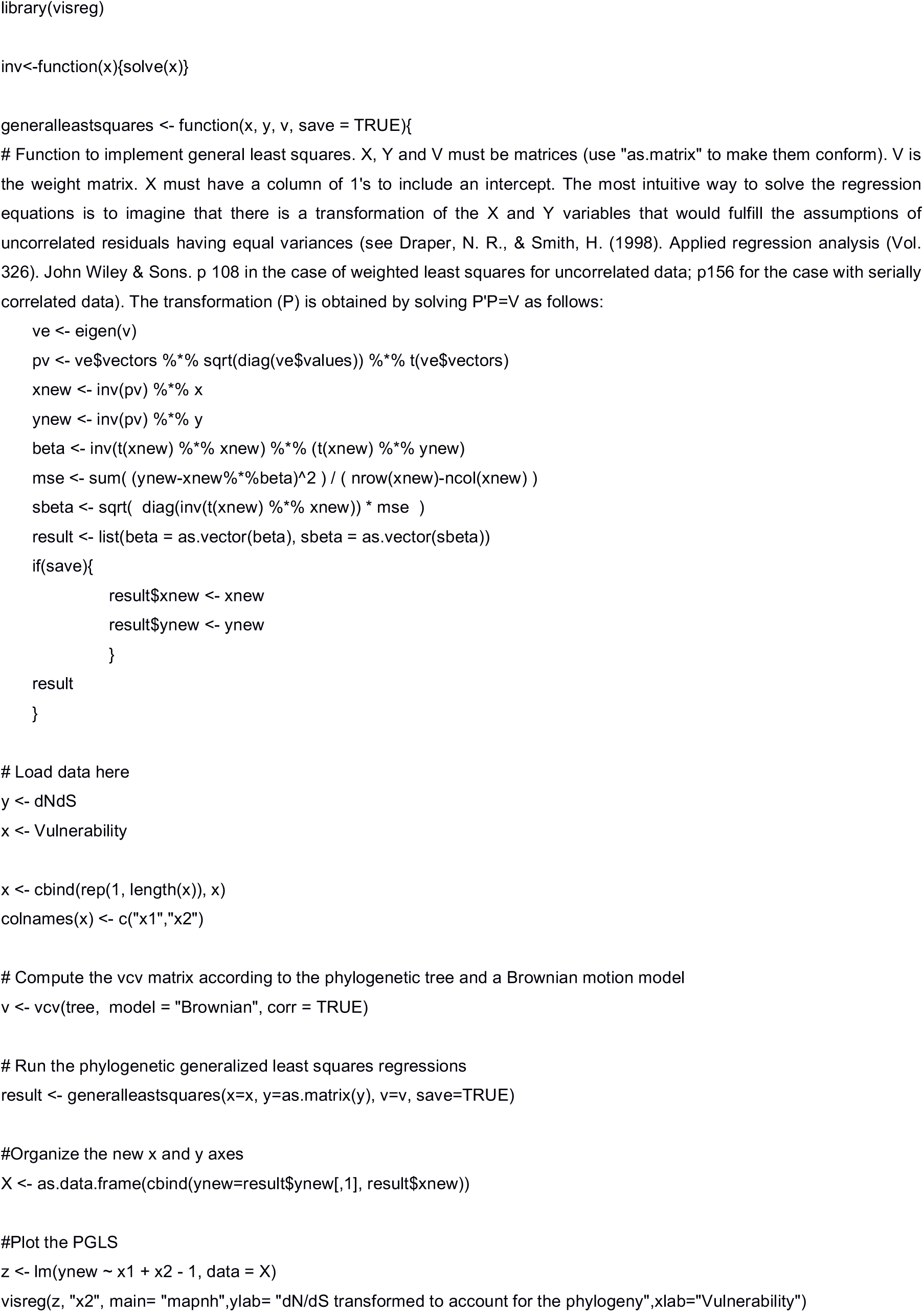
Script permitting to transform x and y axis for accounting for phylogenetic correlation when plotting a phylogenetic generalized least squares regression.

**Supplementary Material S7.**
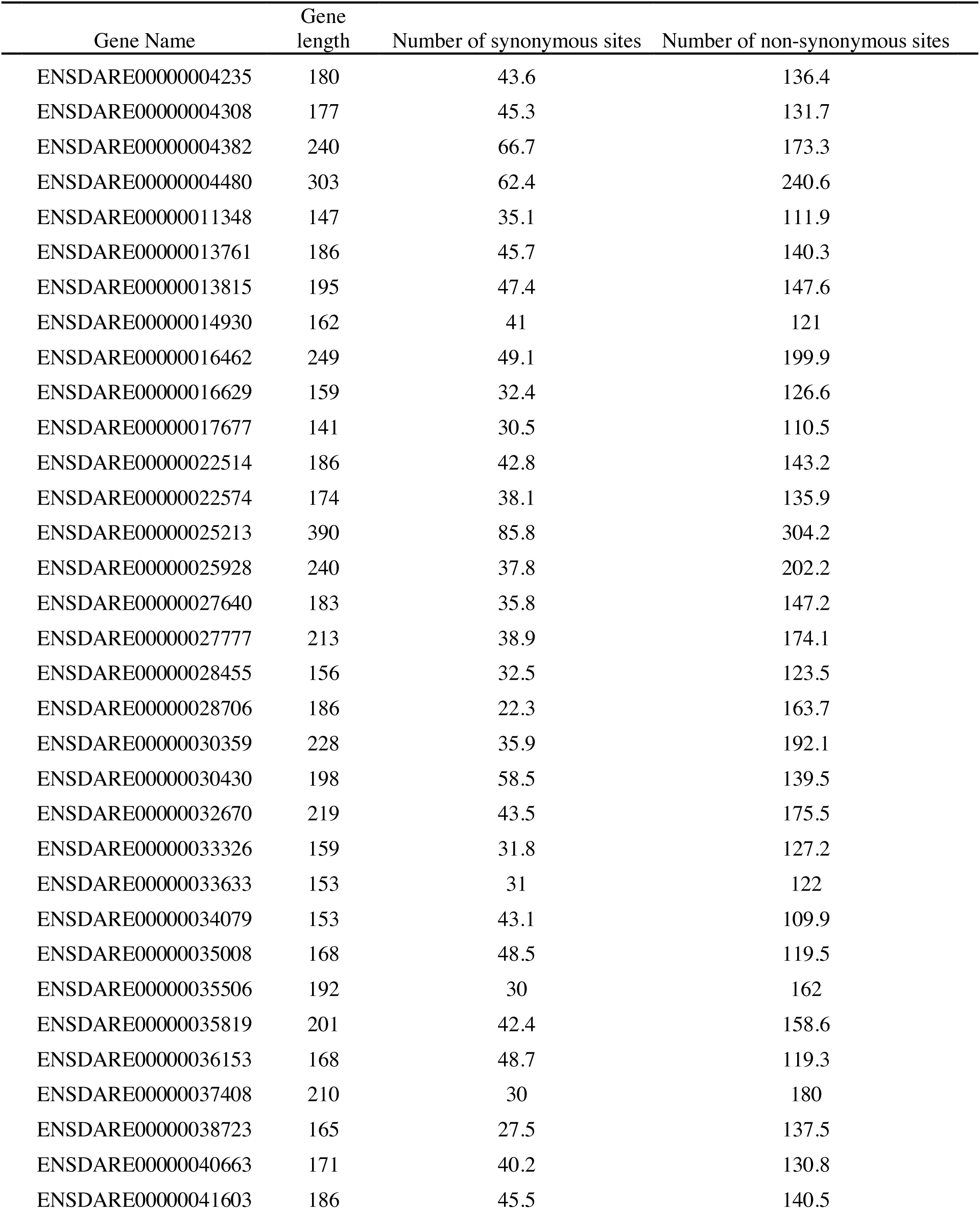
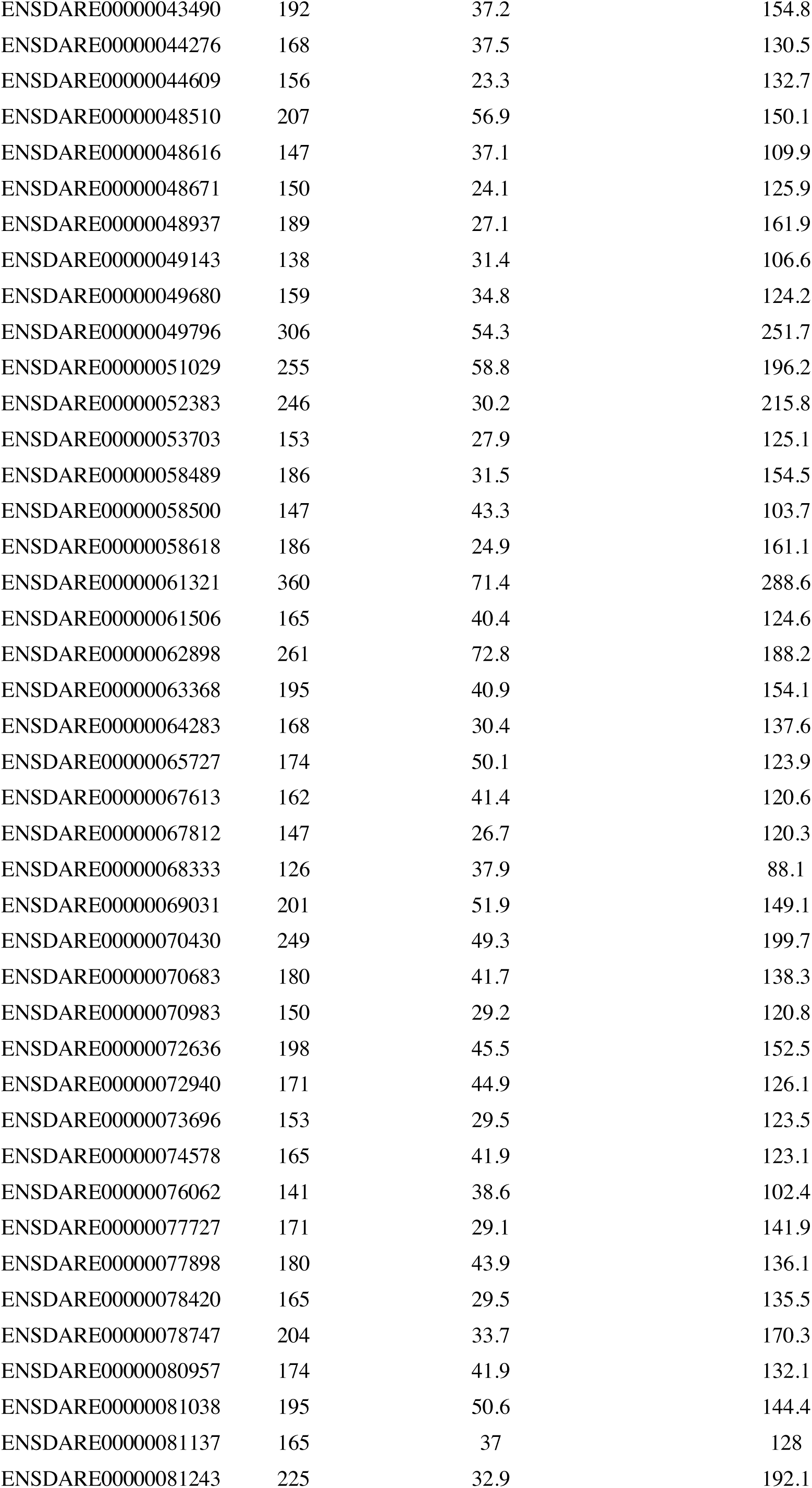
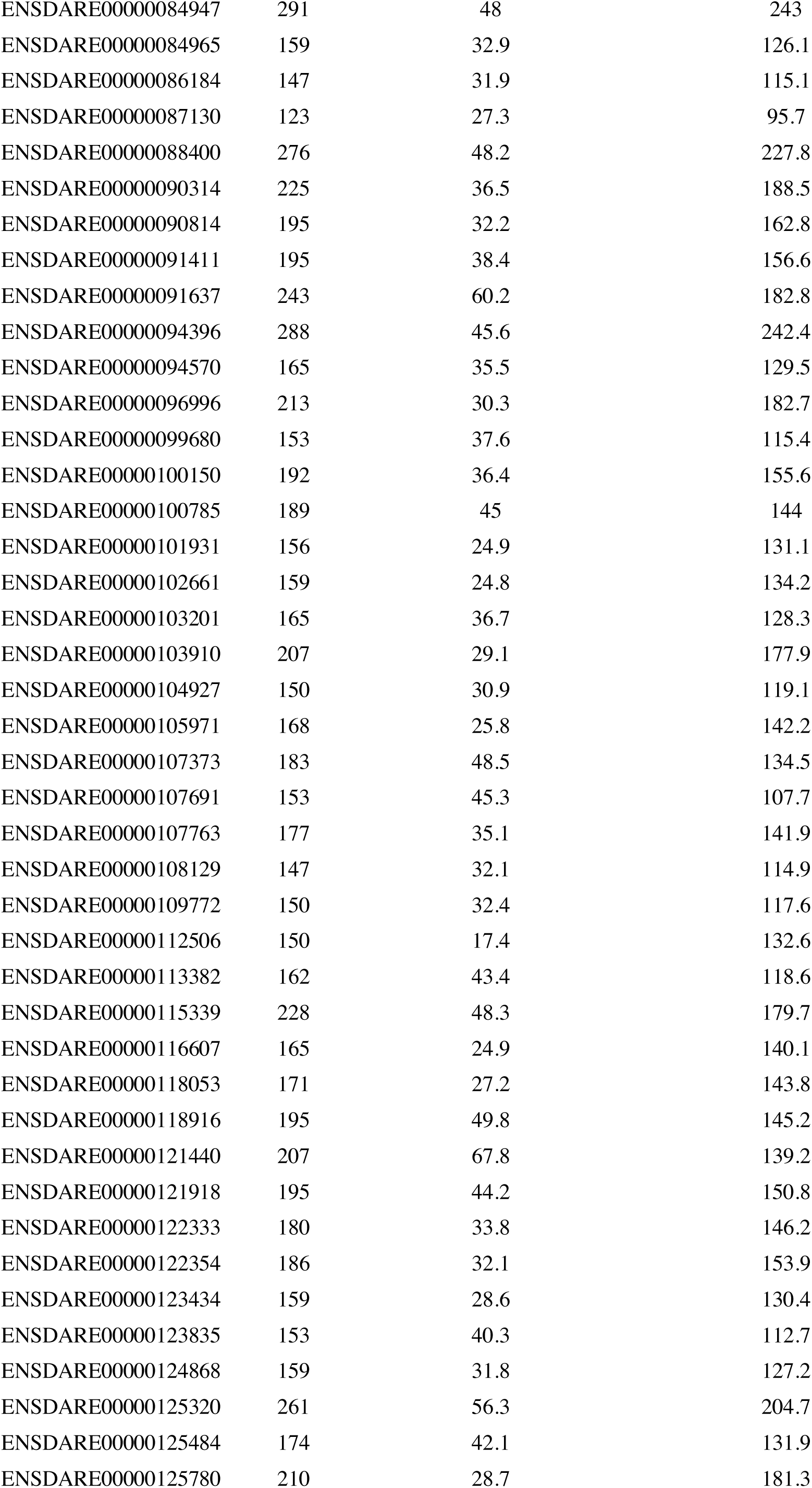
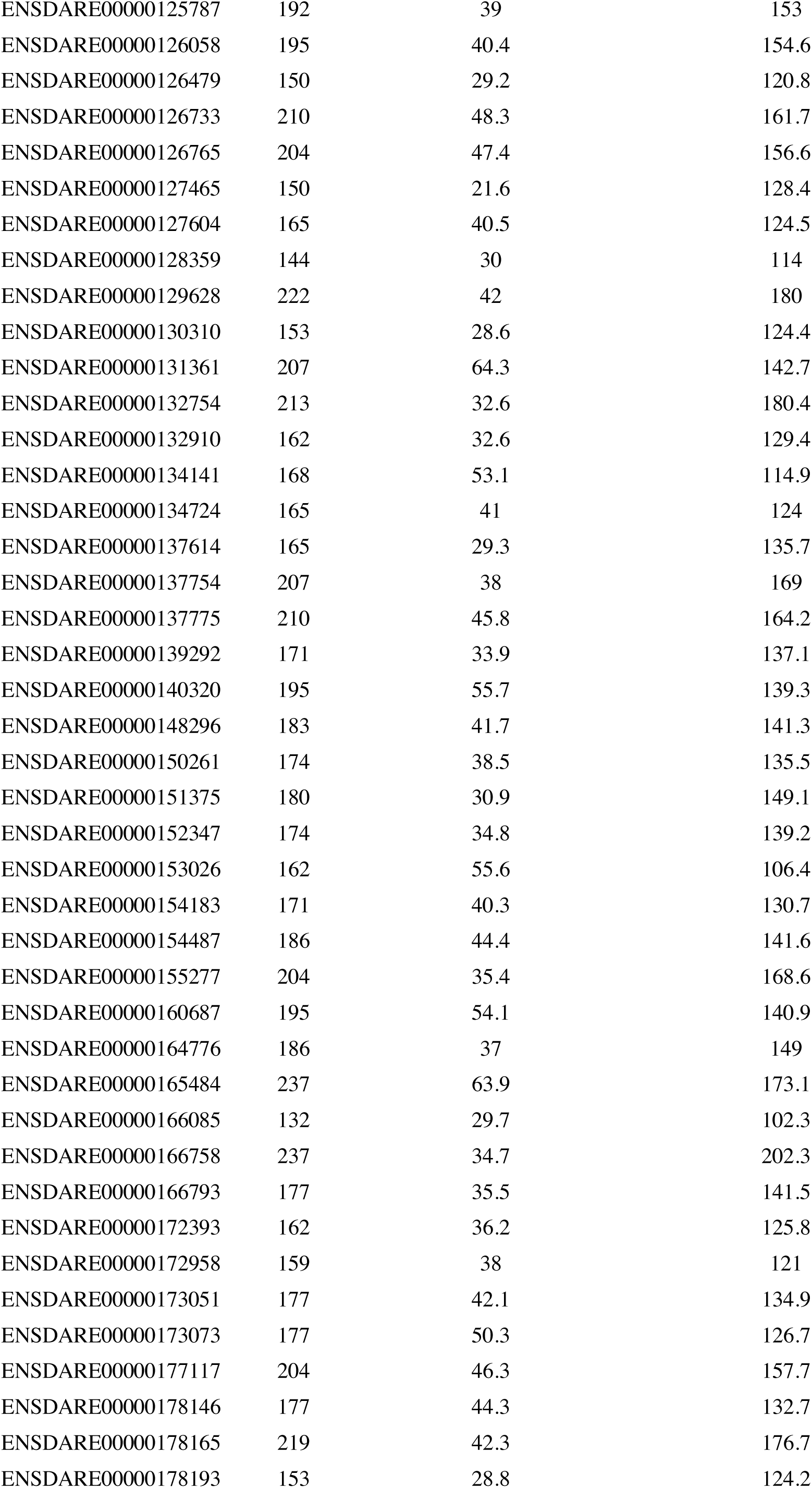
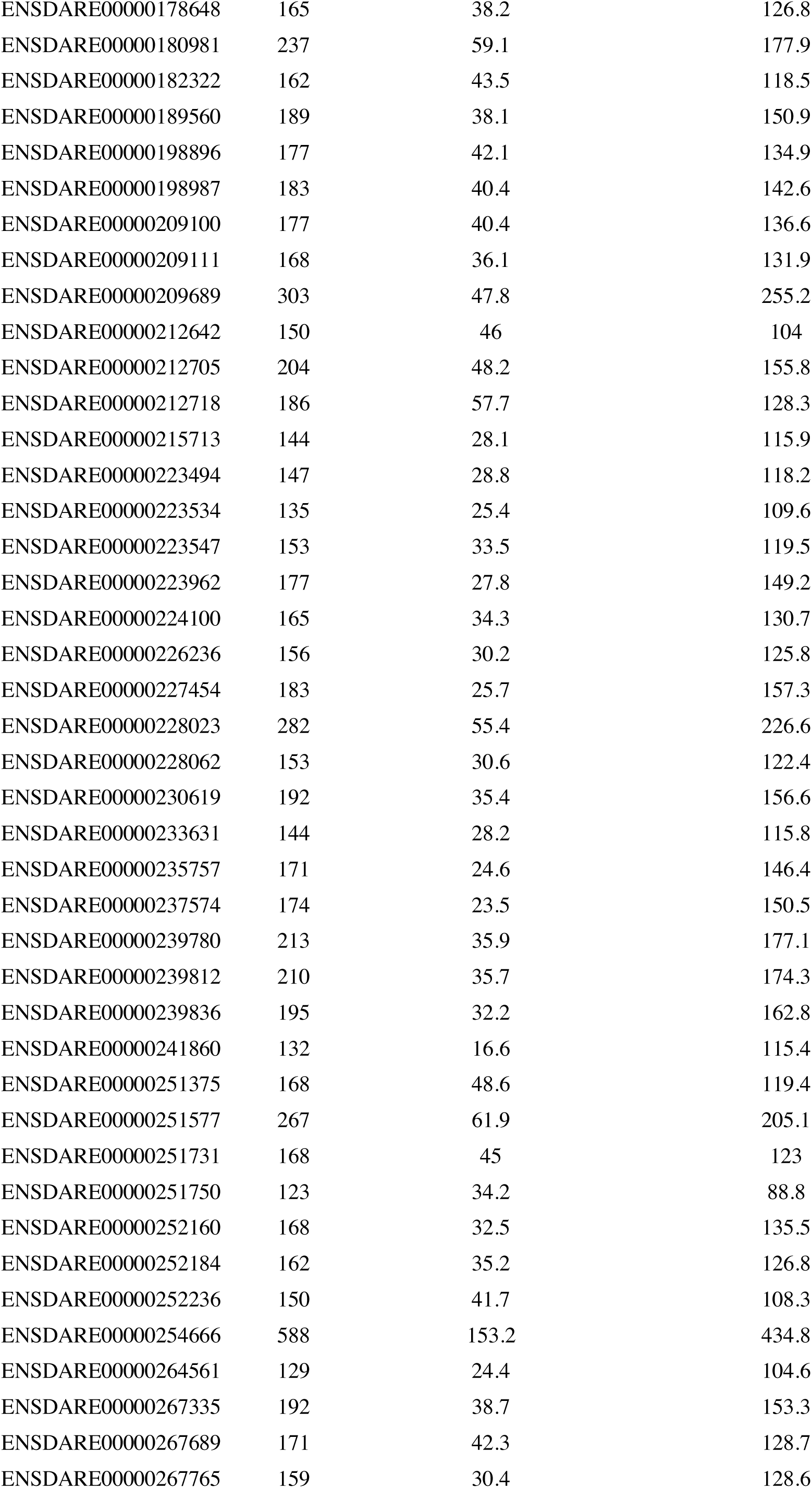
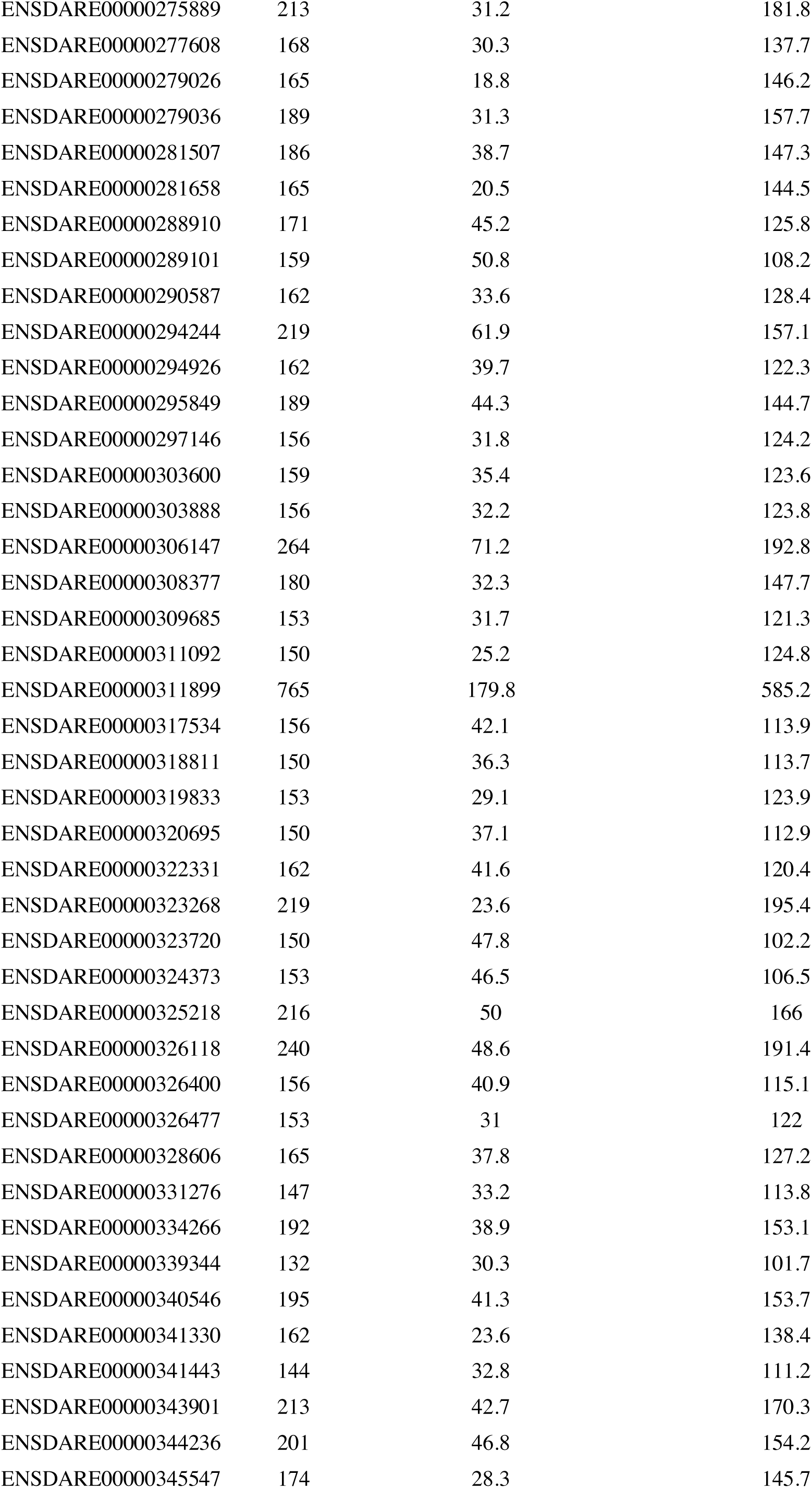
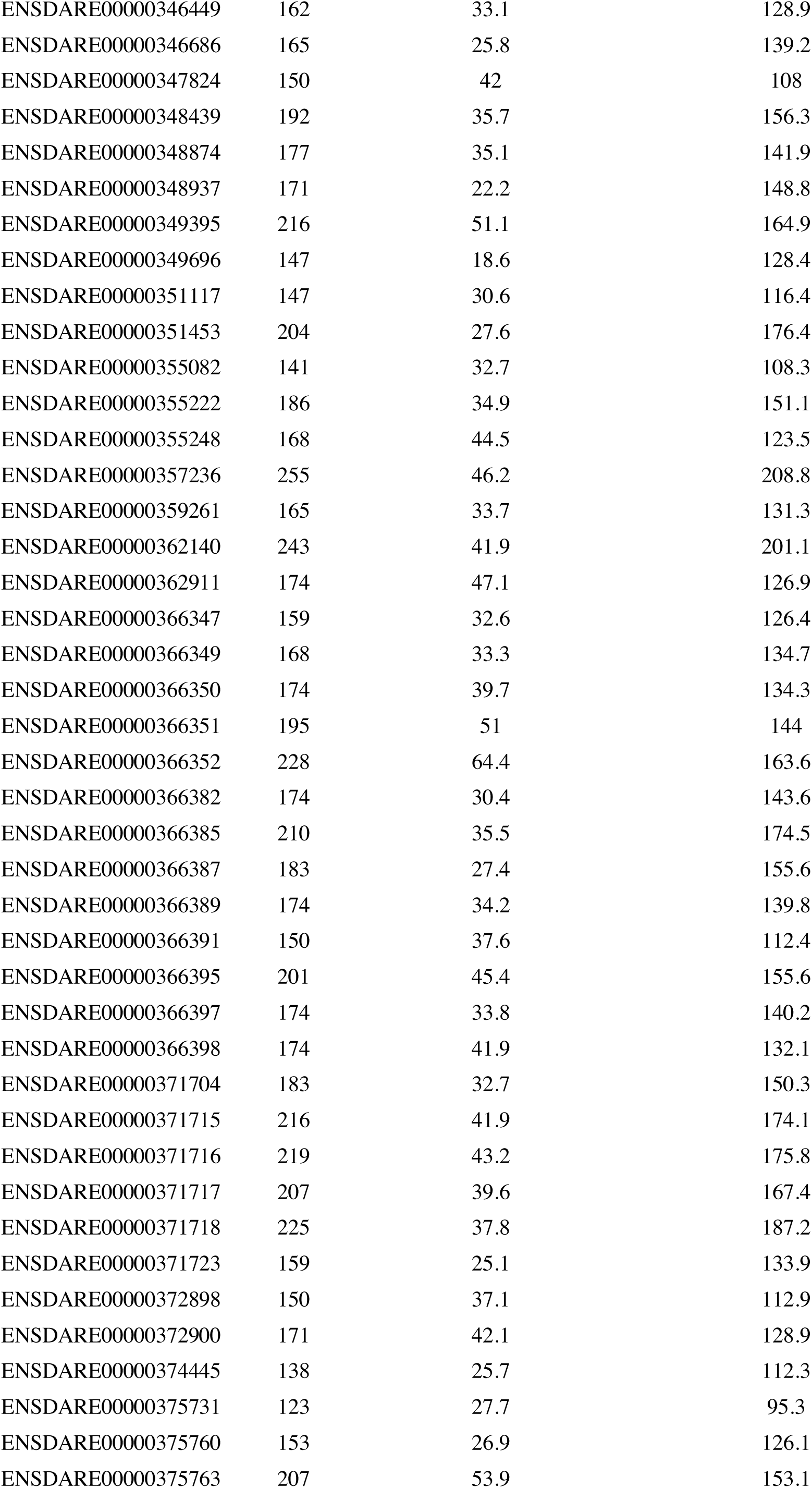
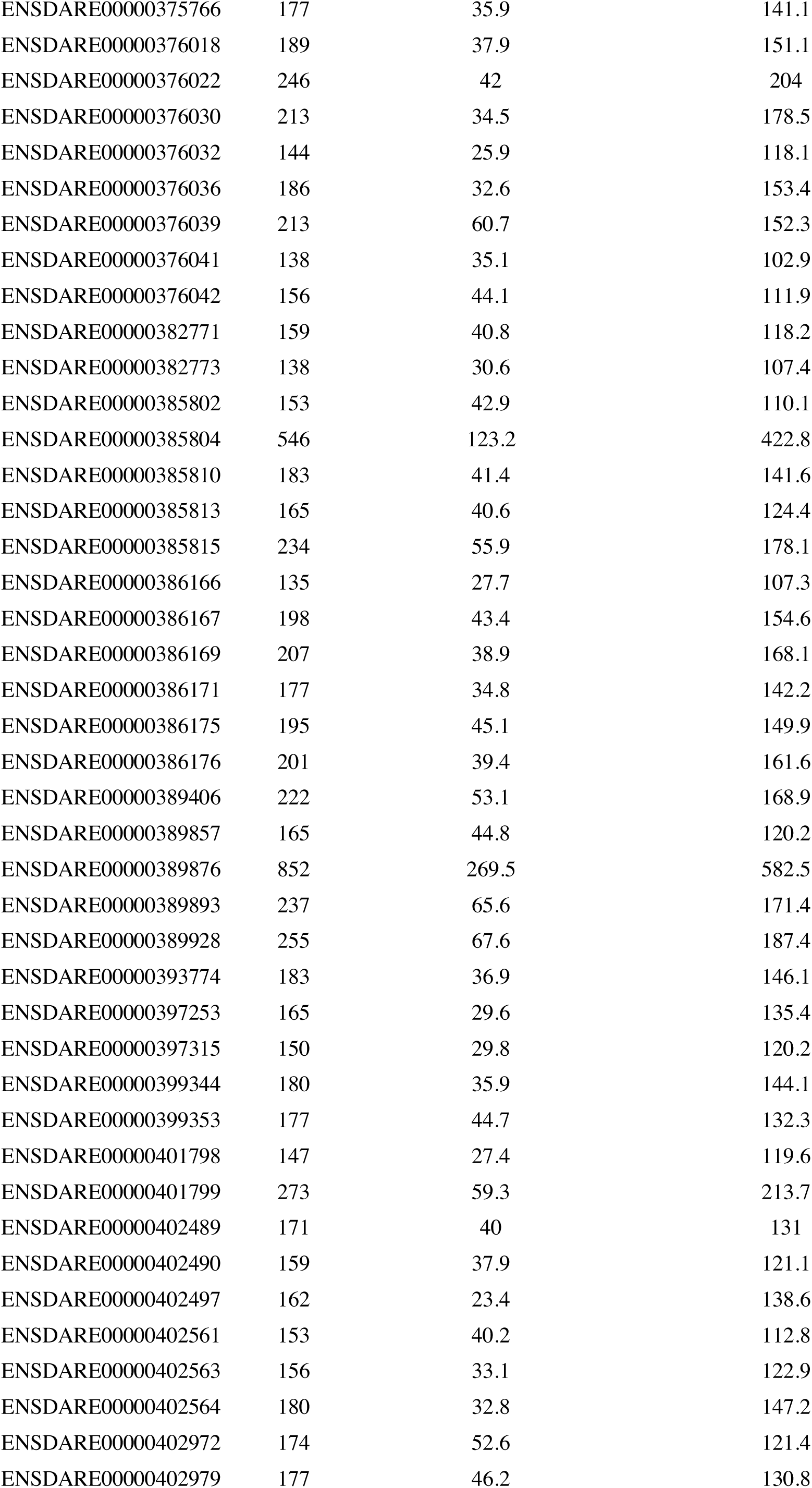
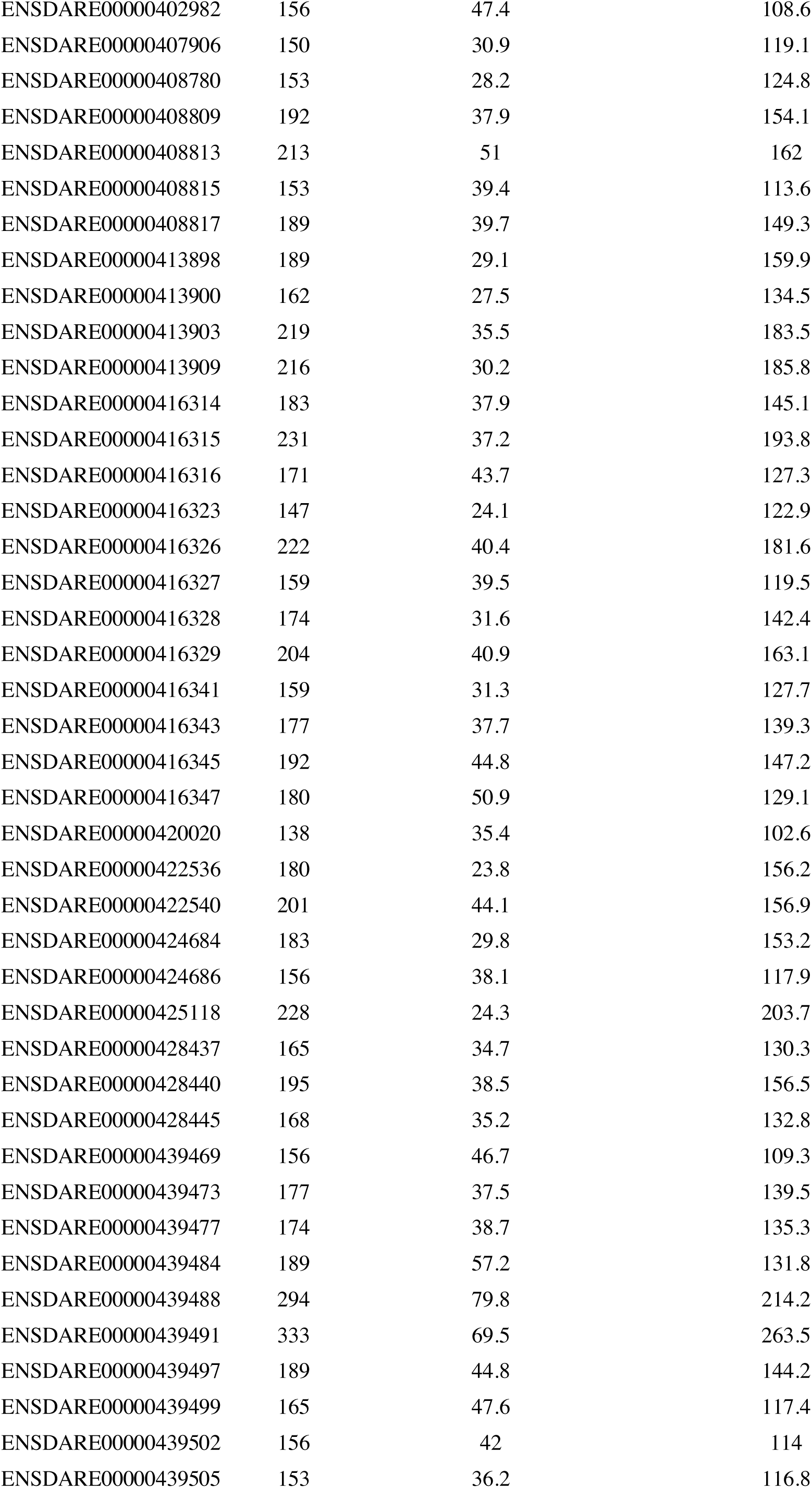
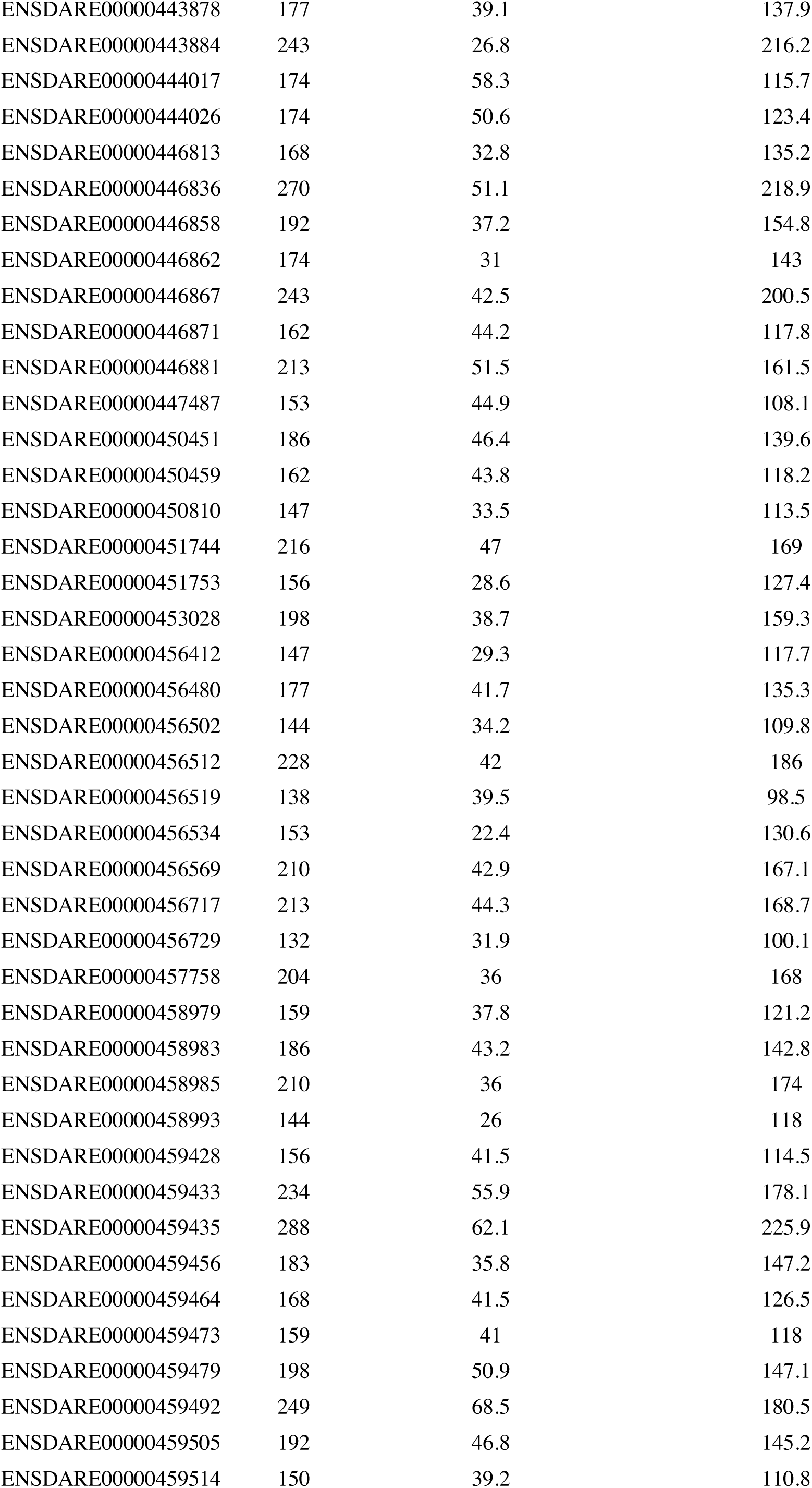
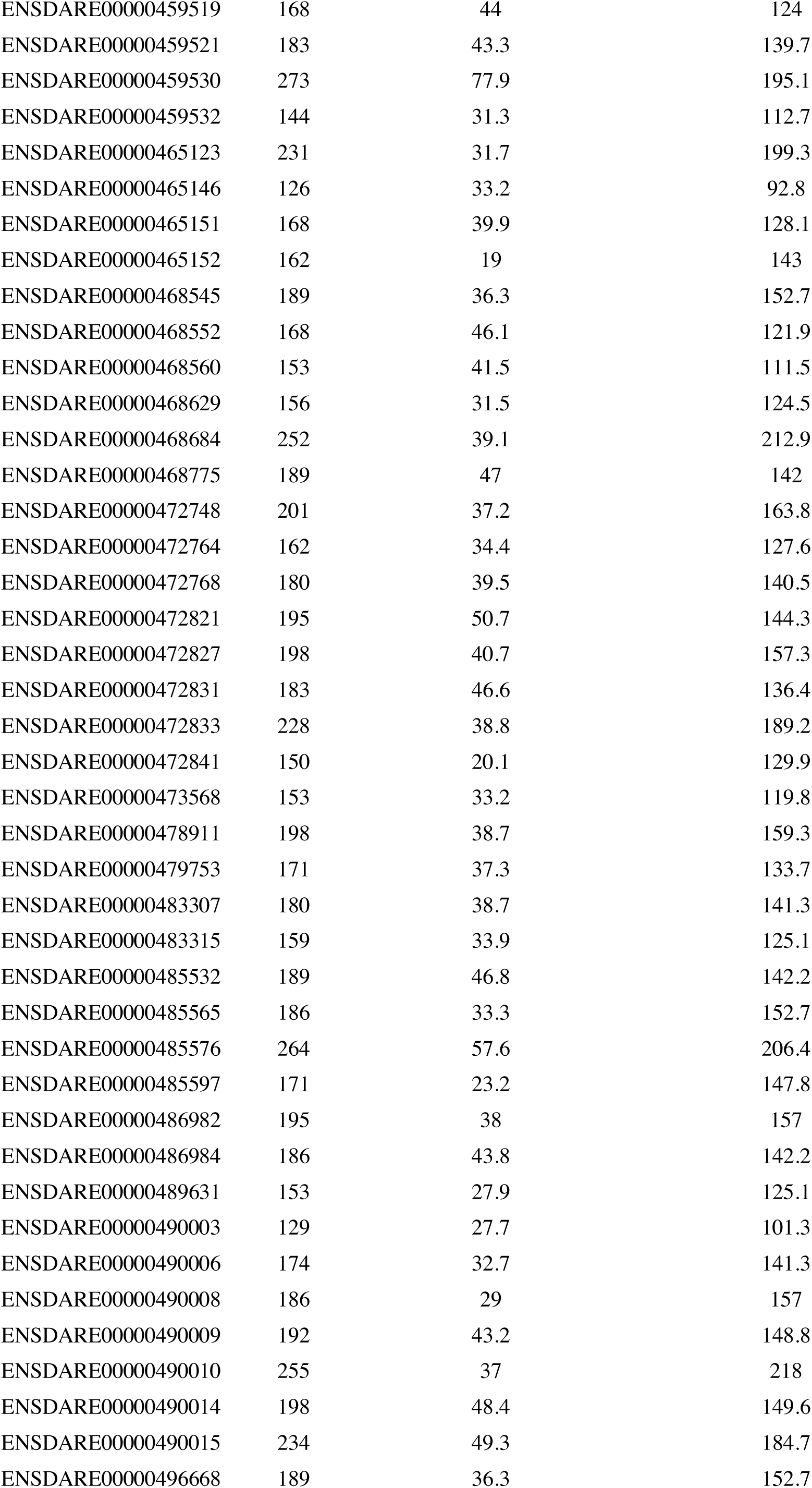
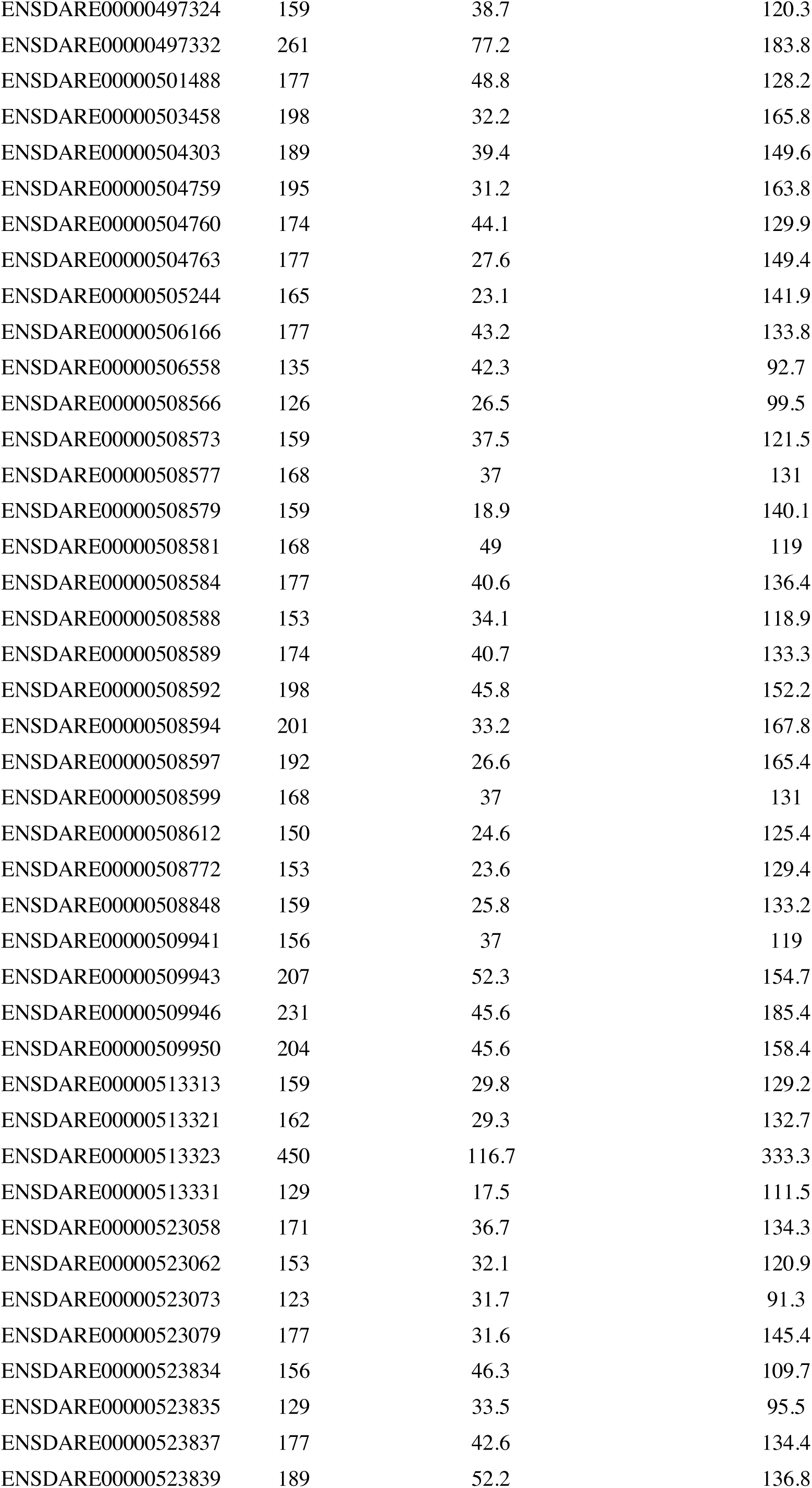
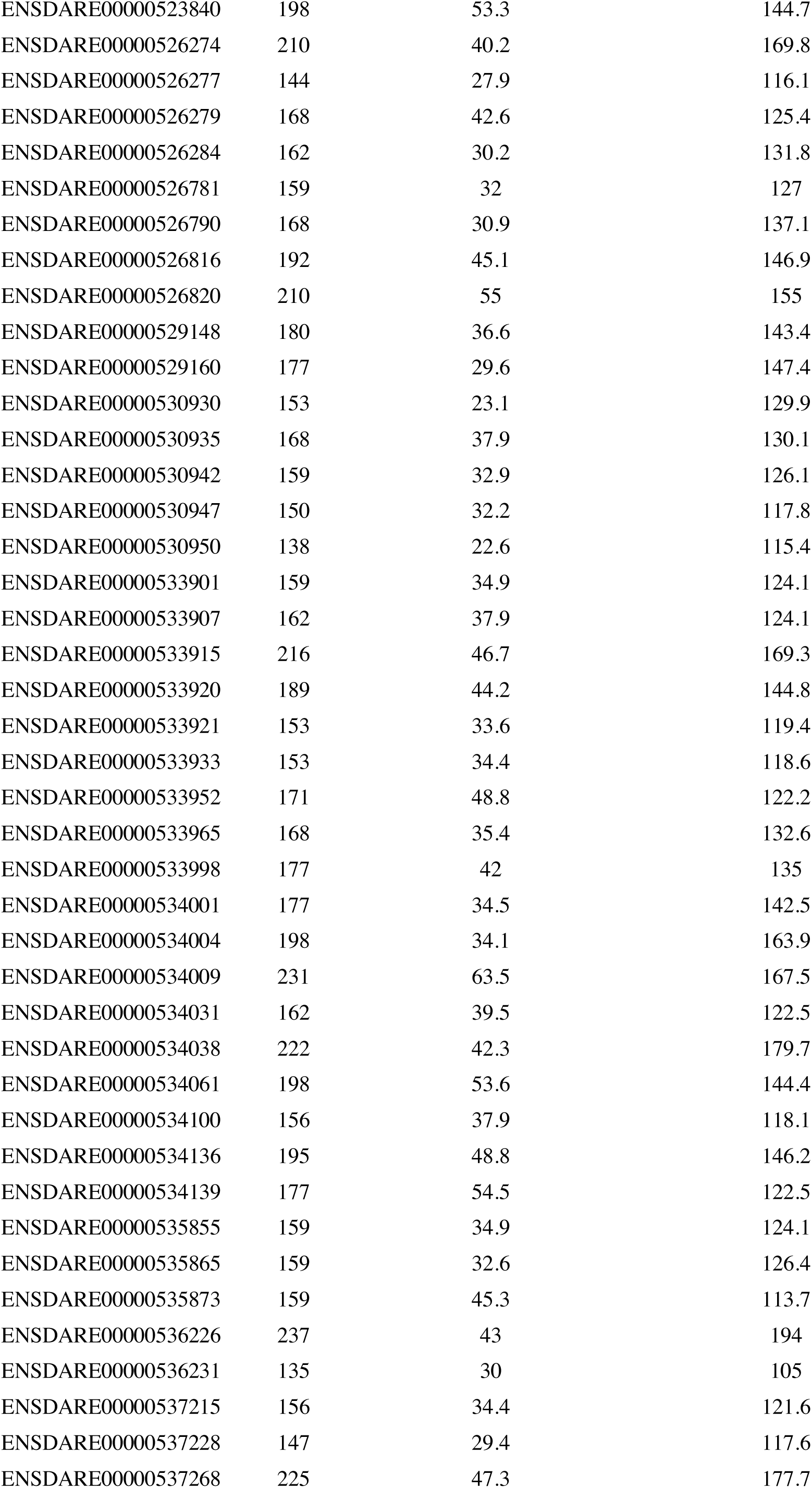
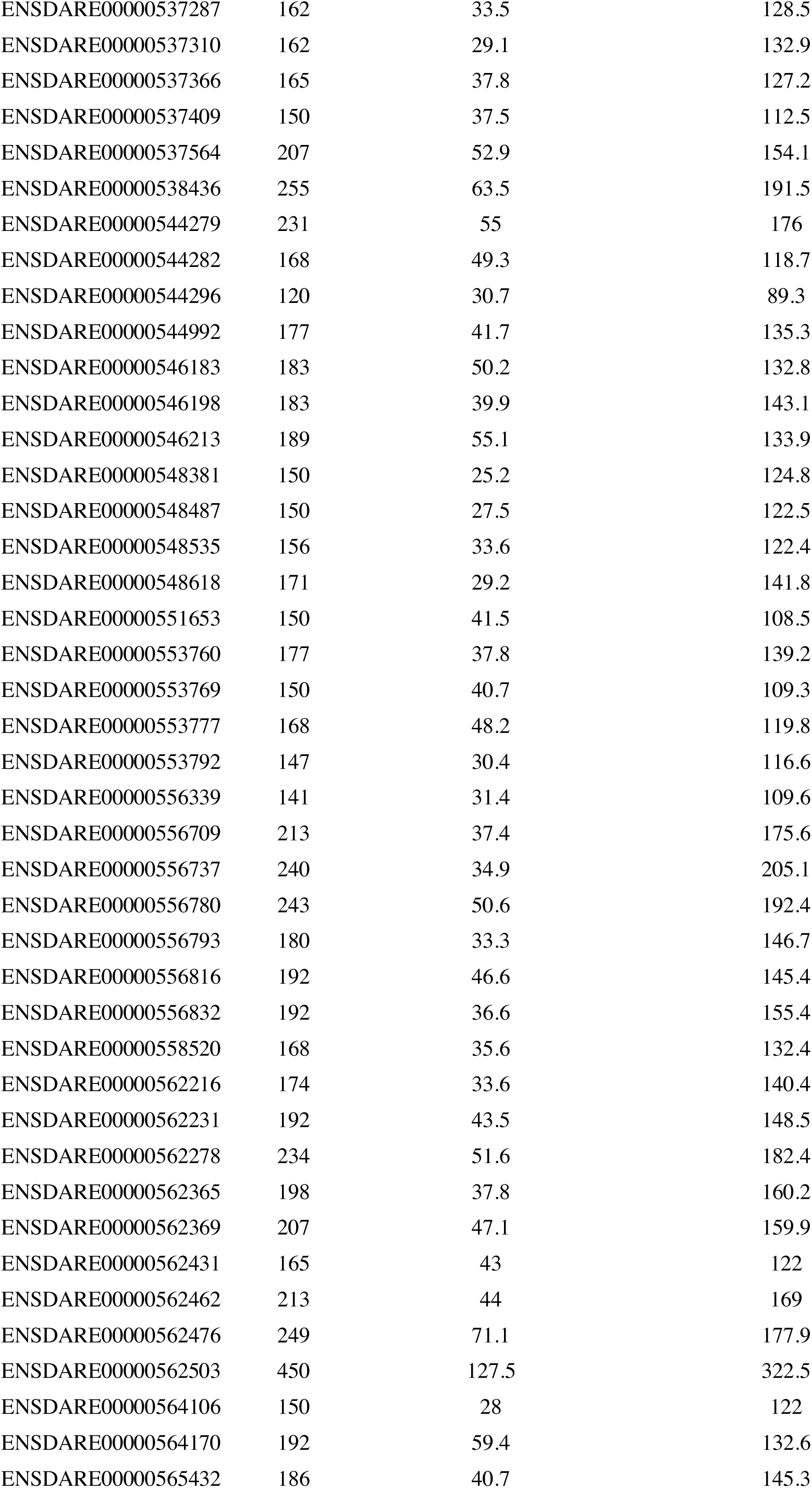
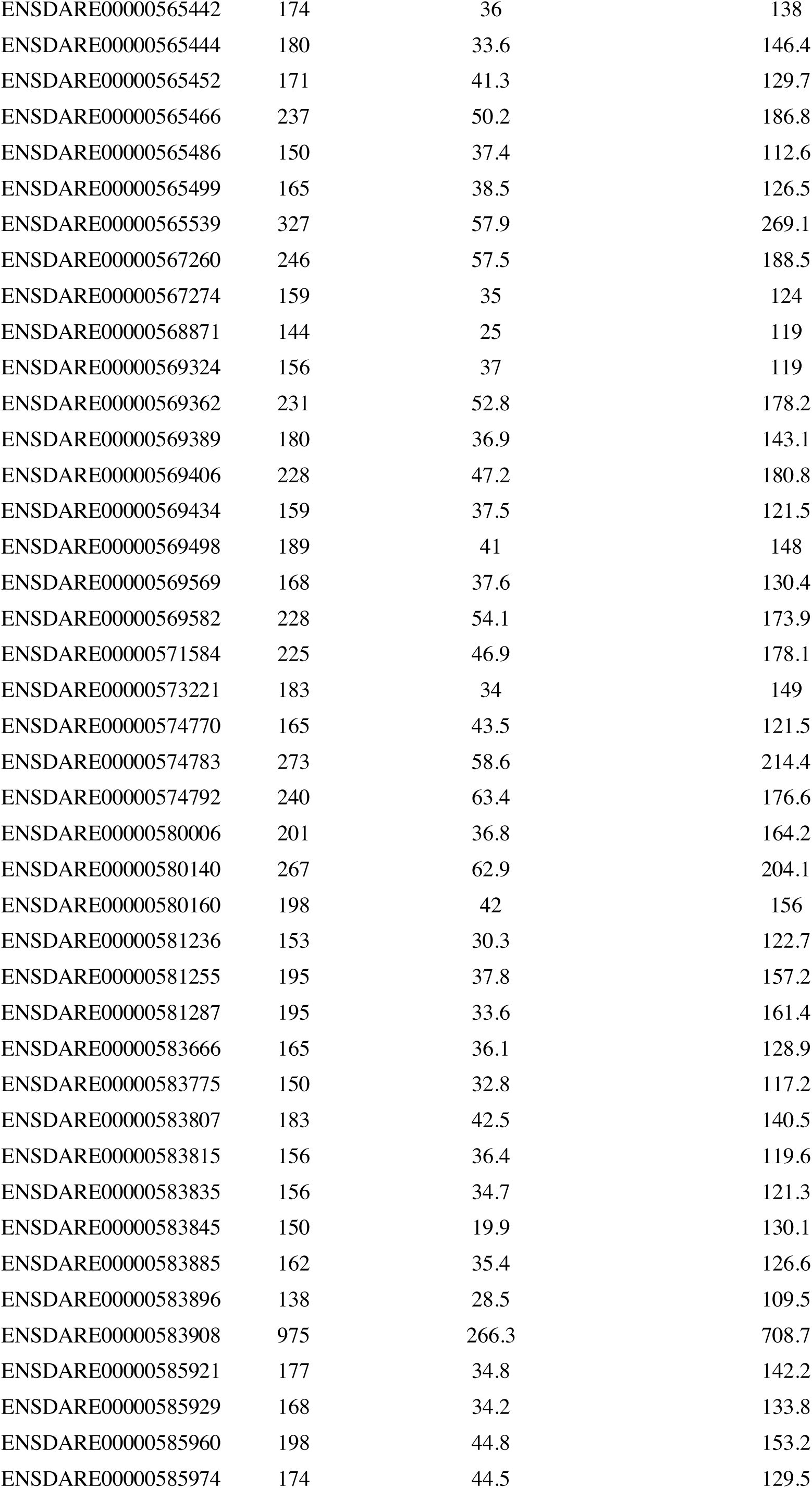
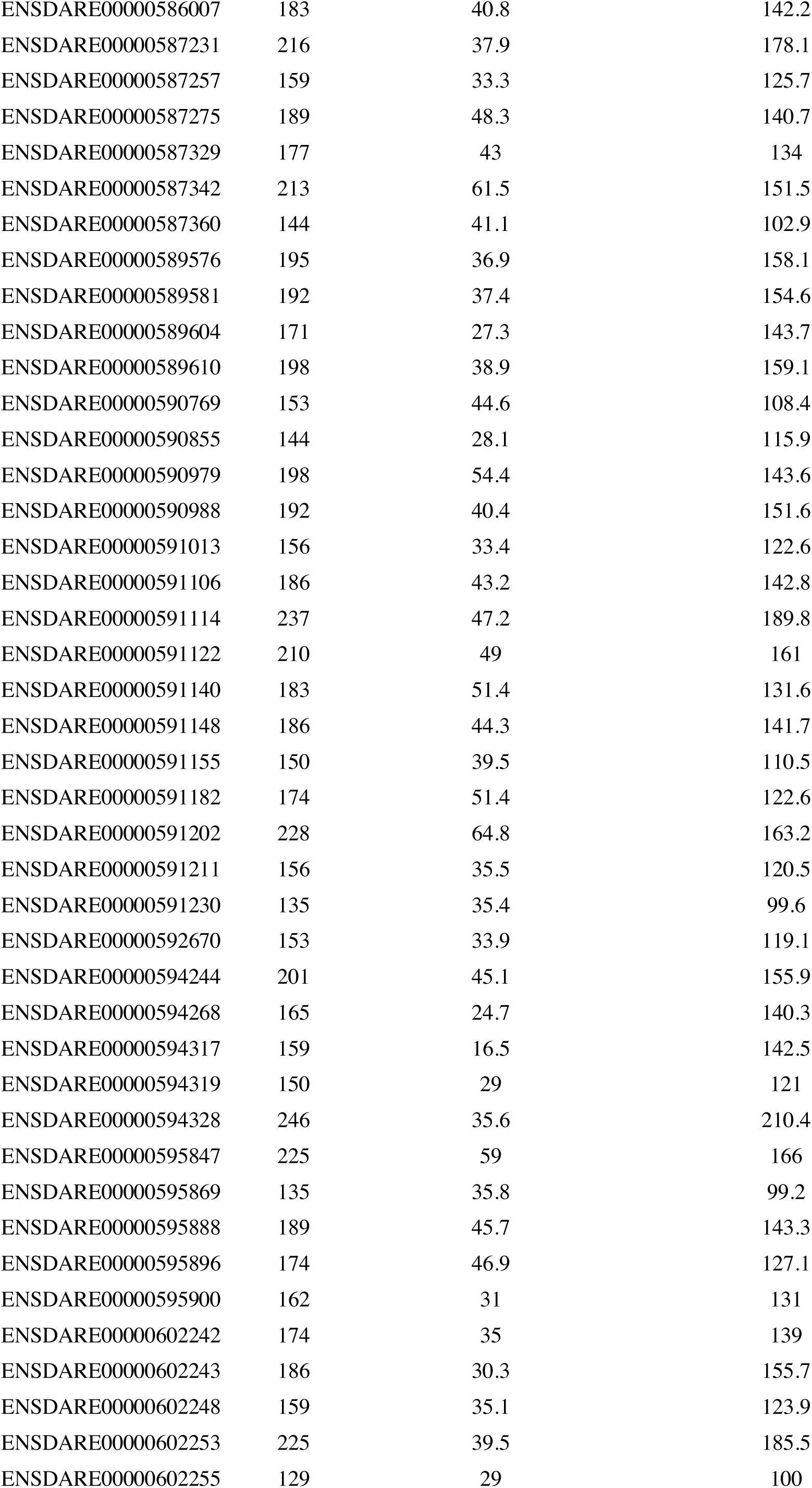
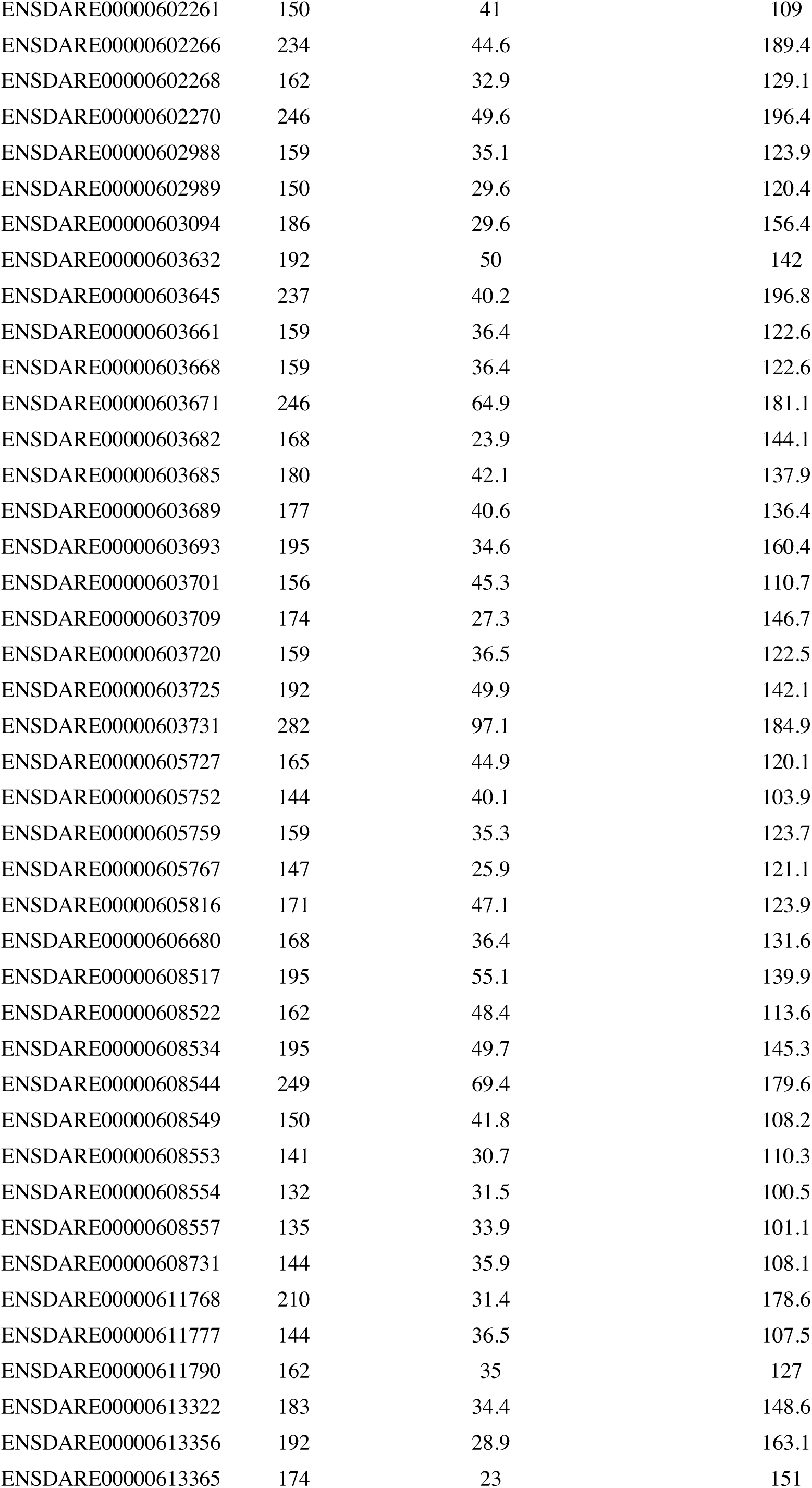
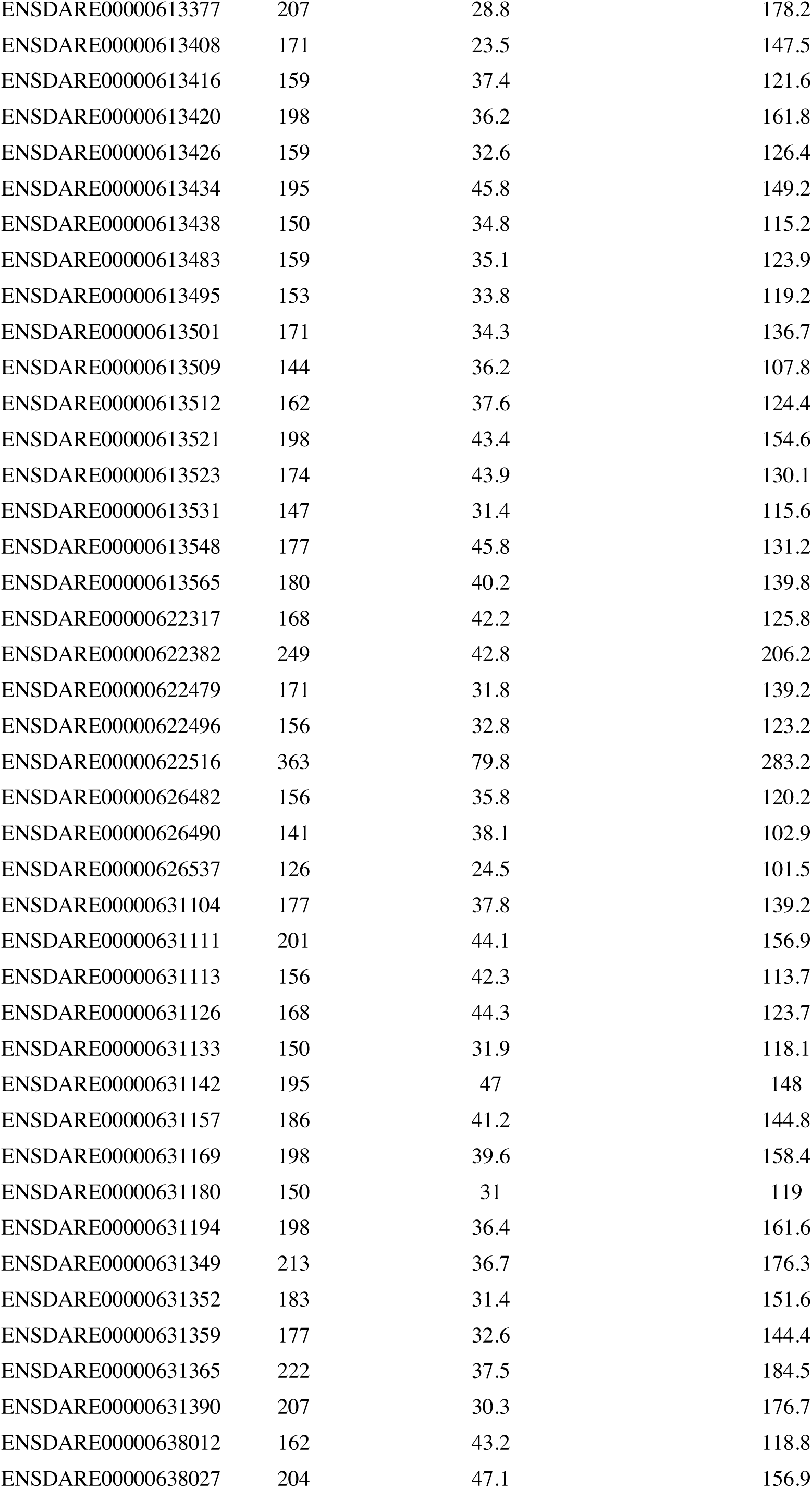
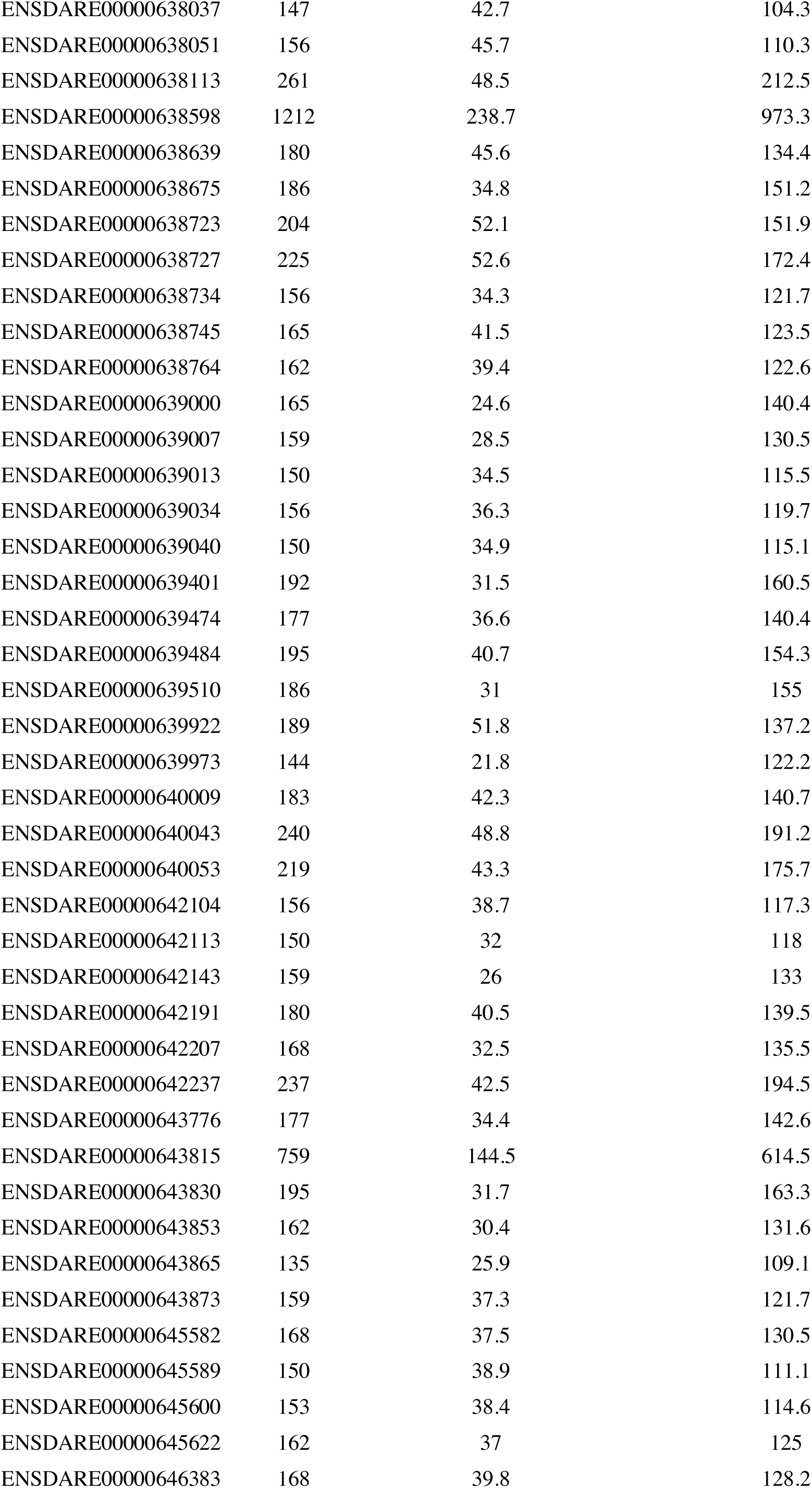
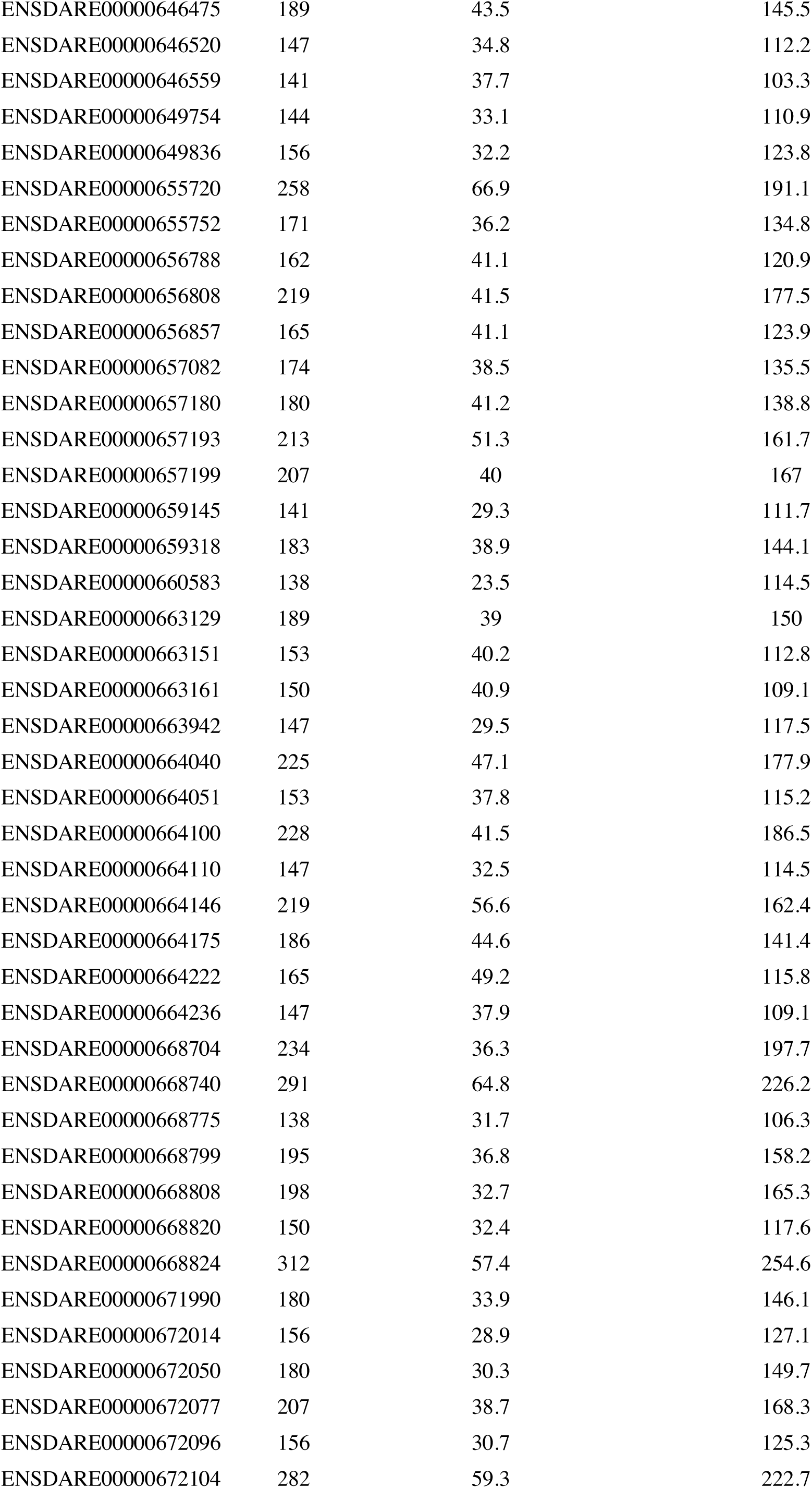
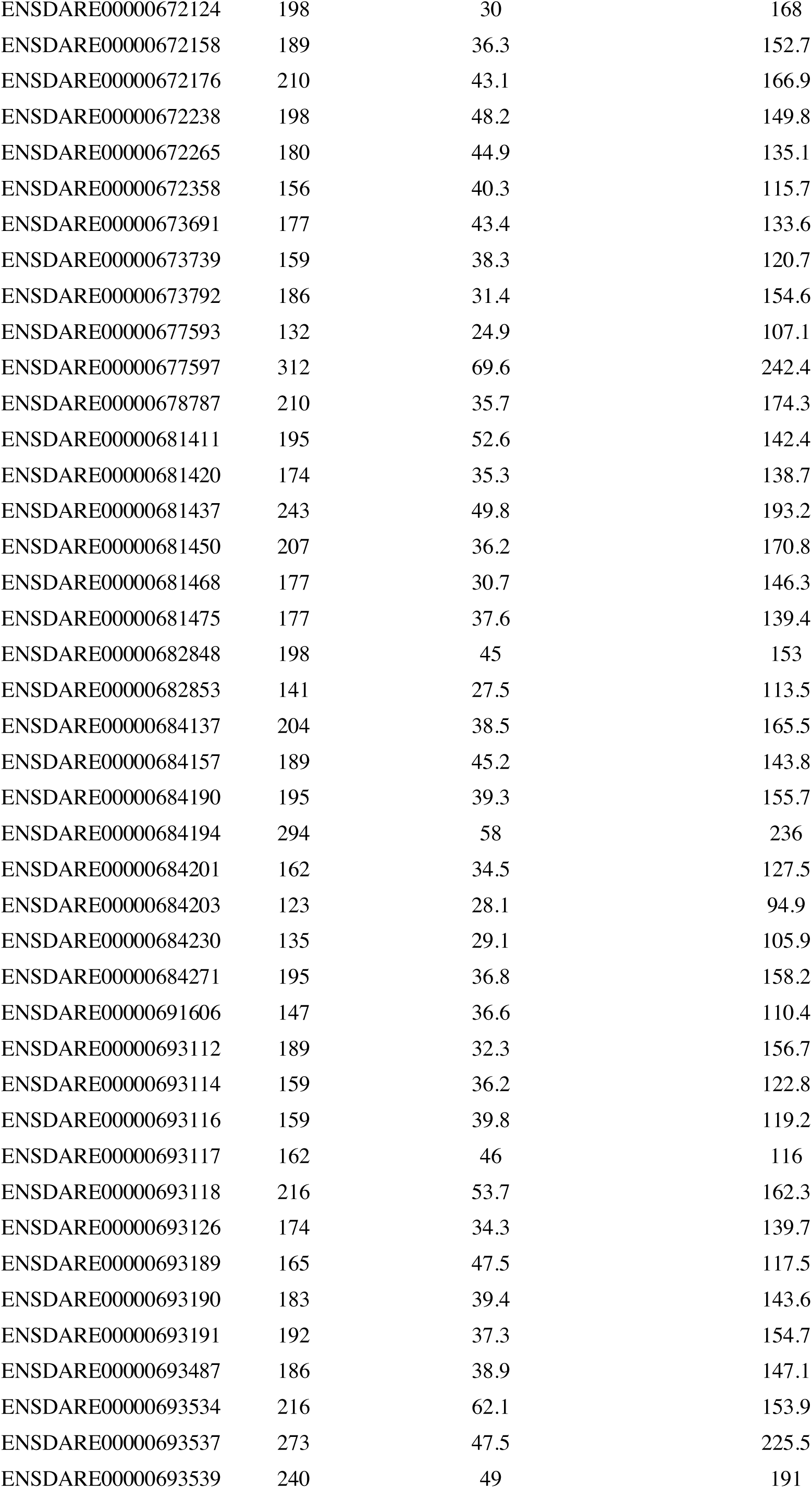
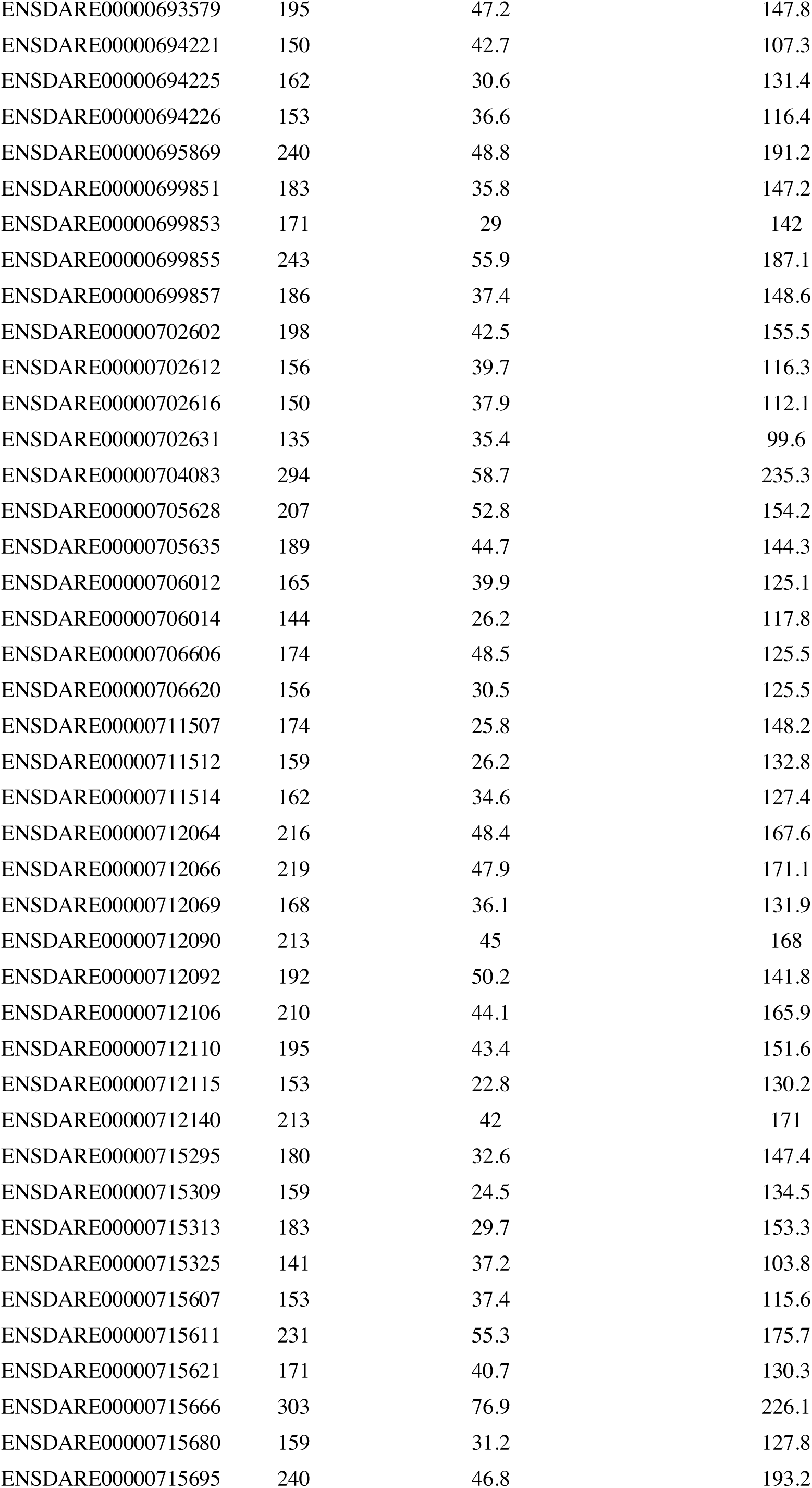
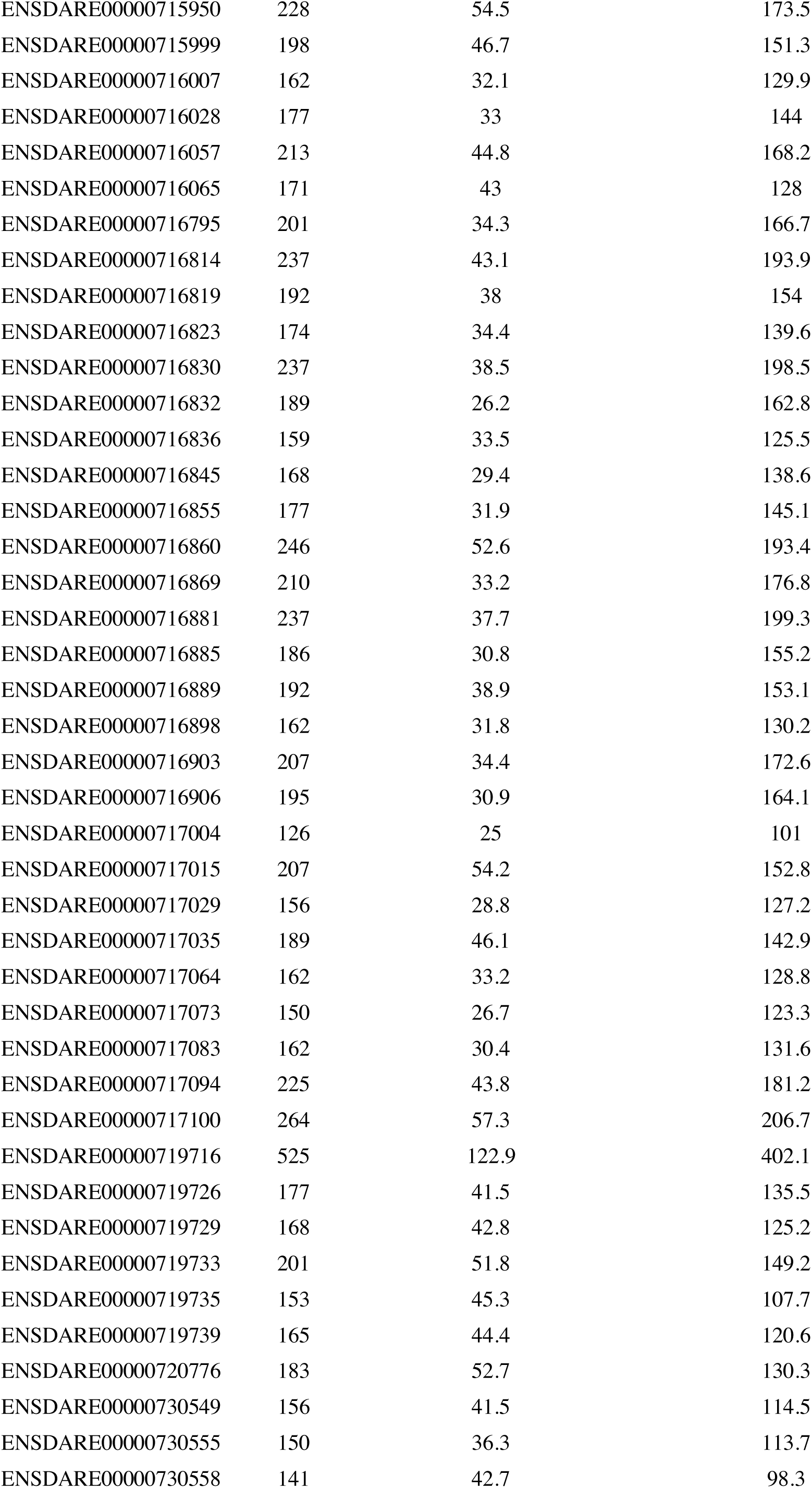
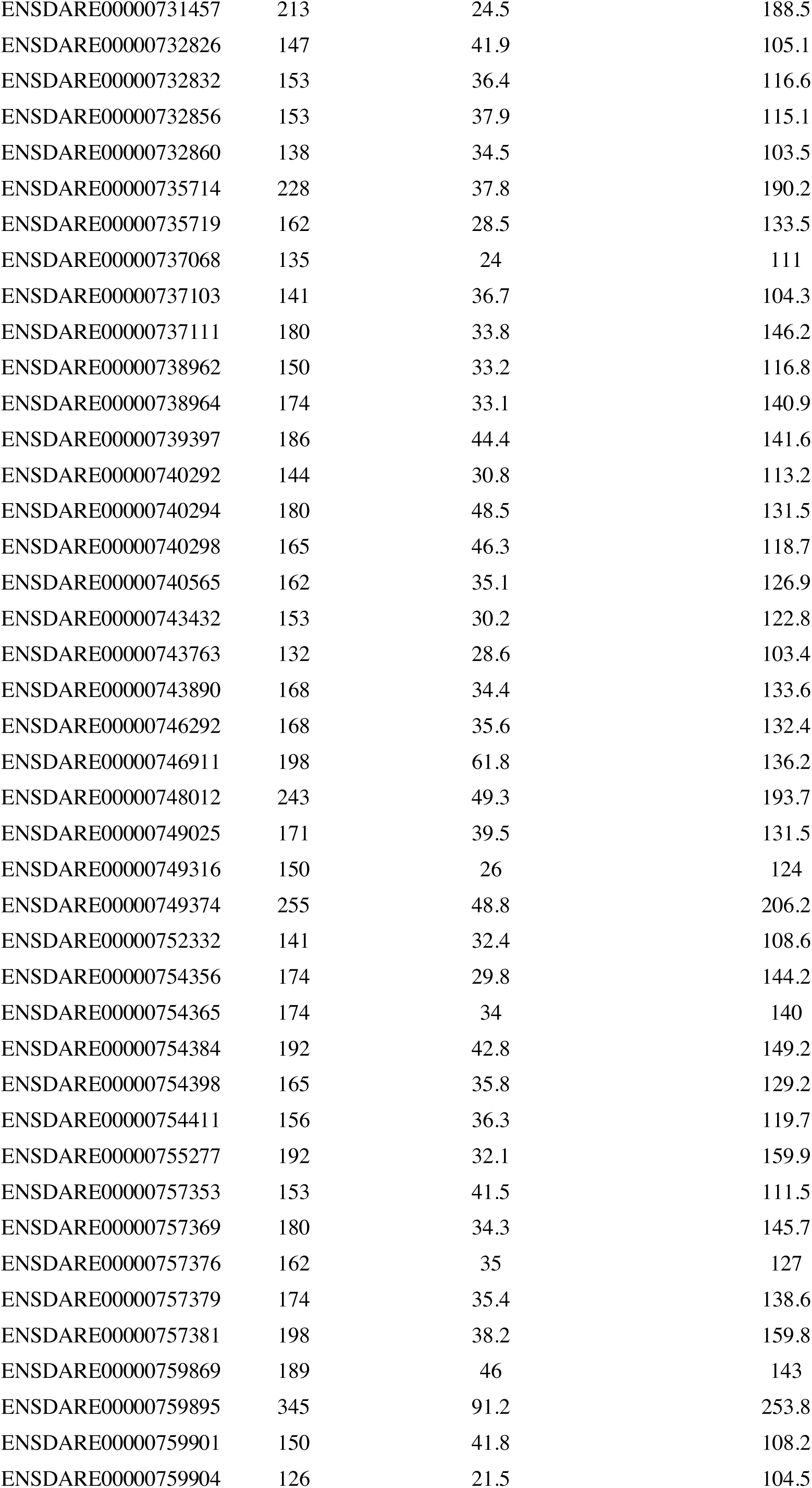
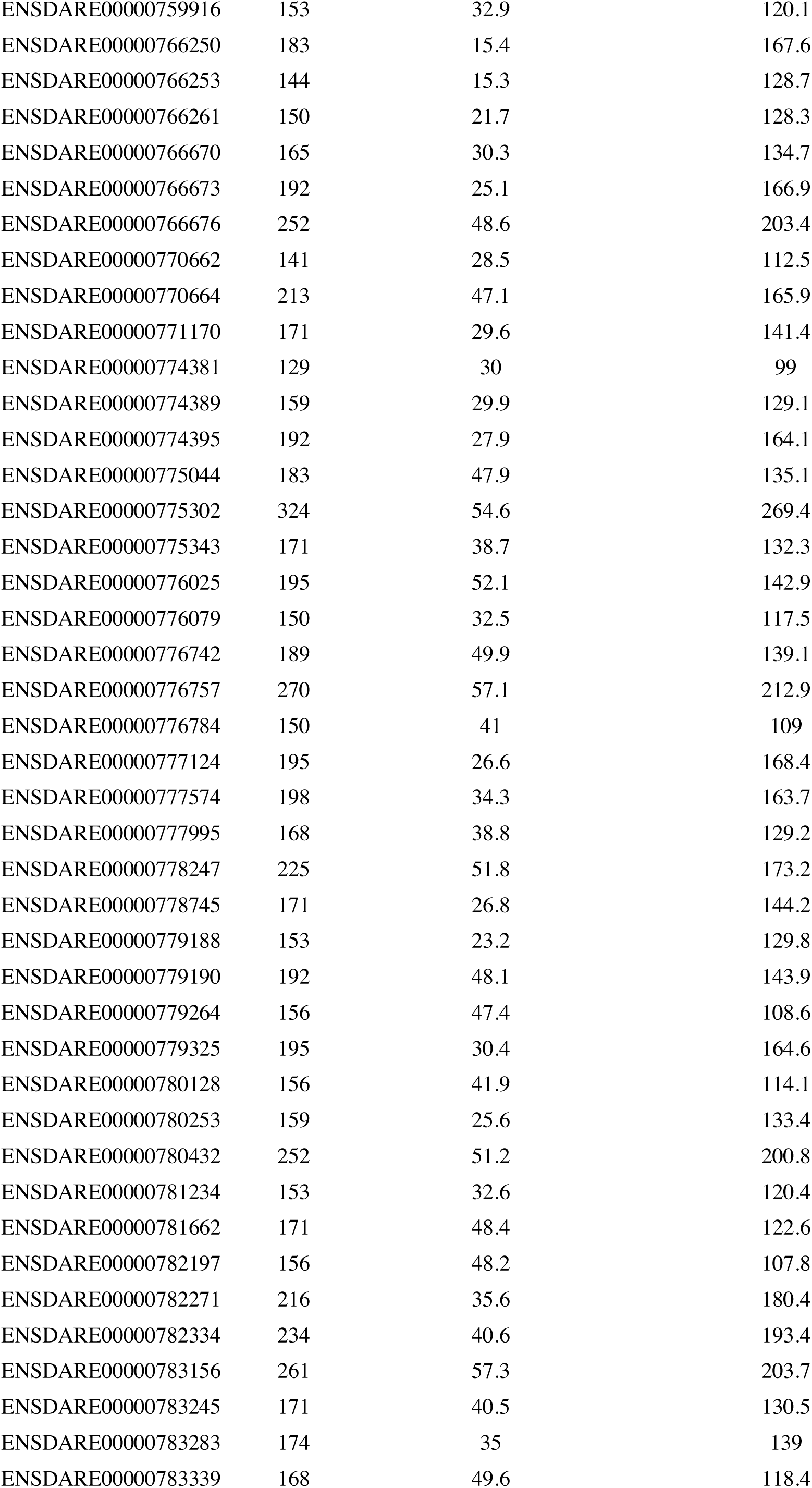
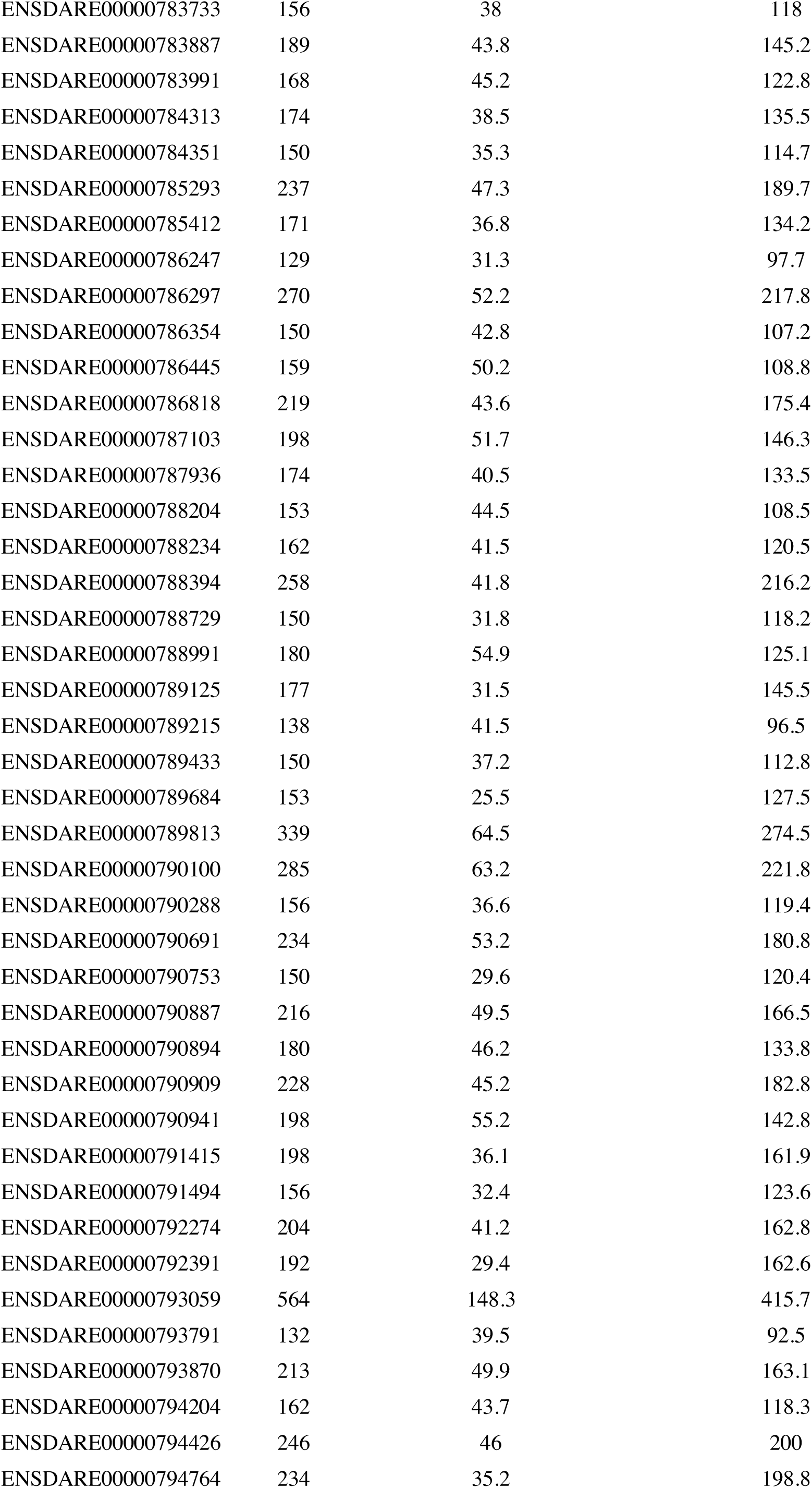
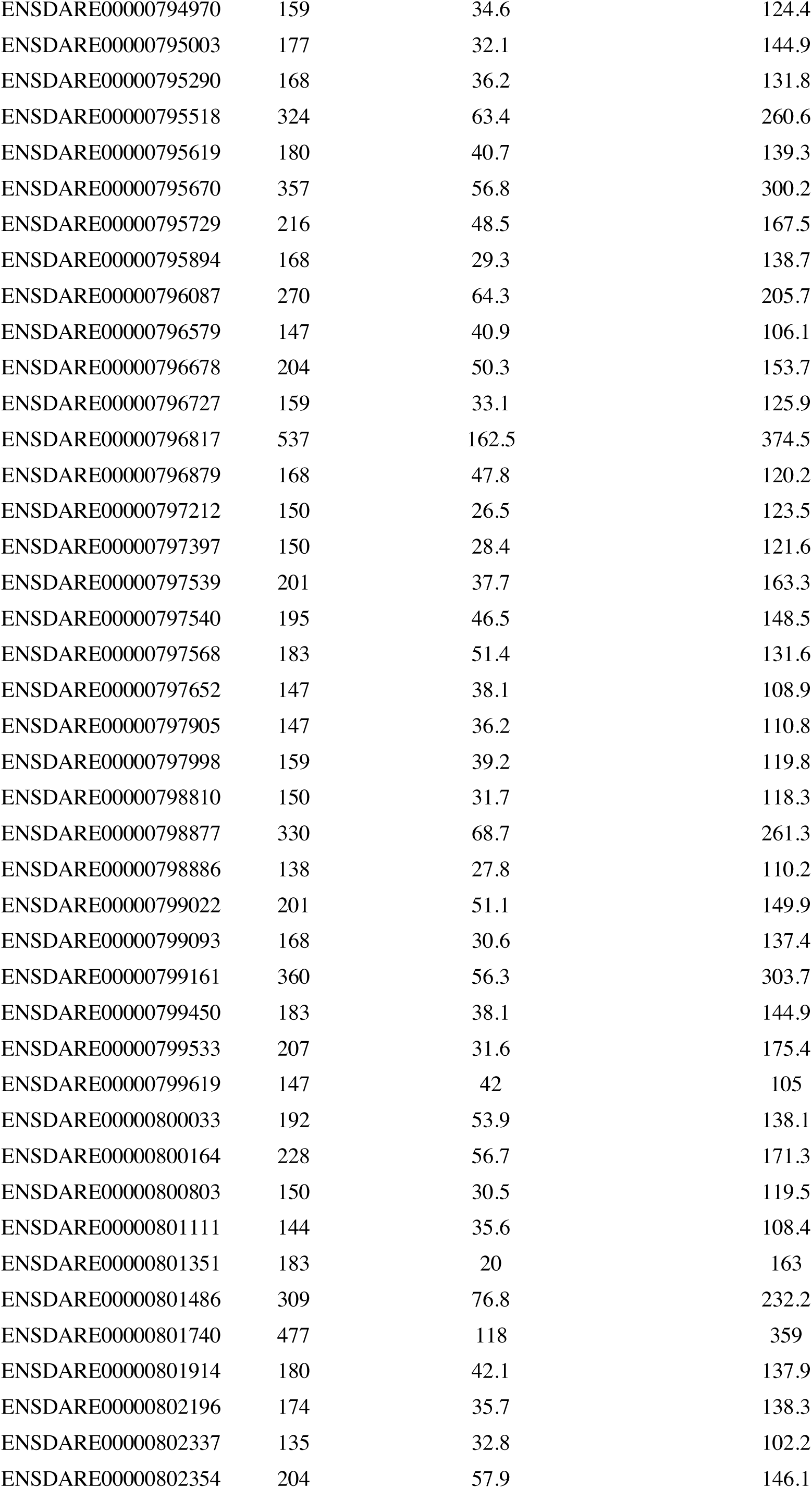
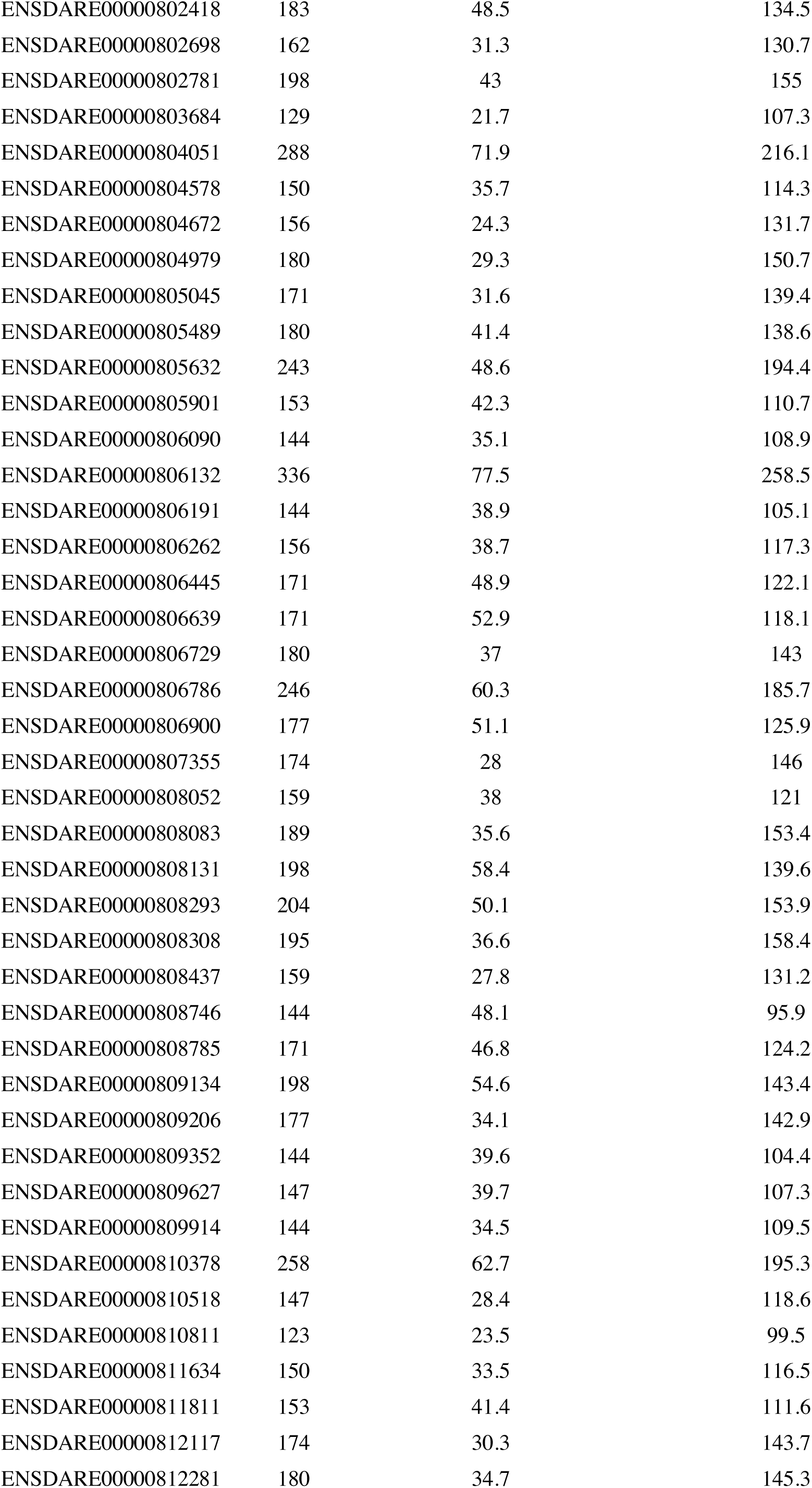
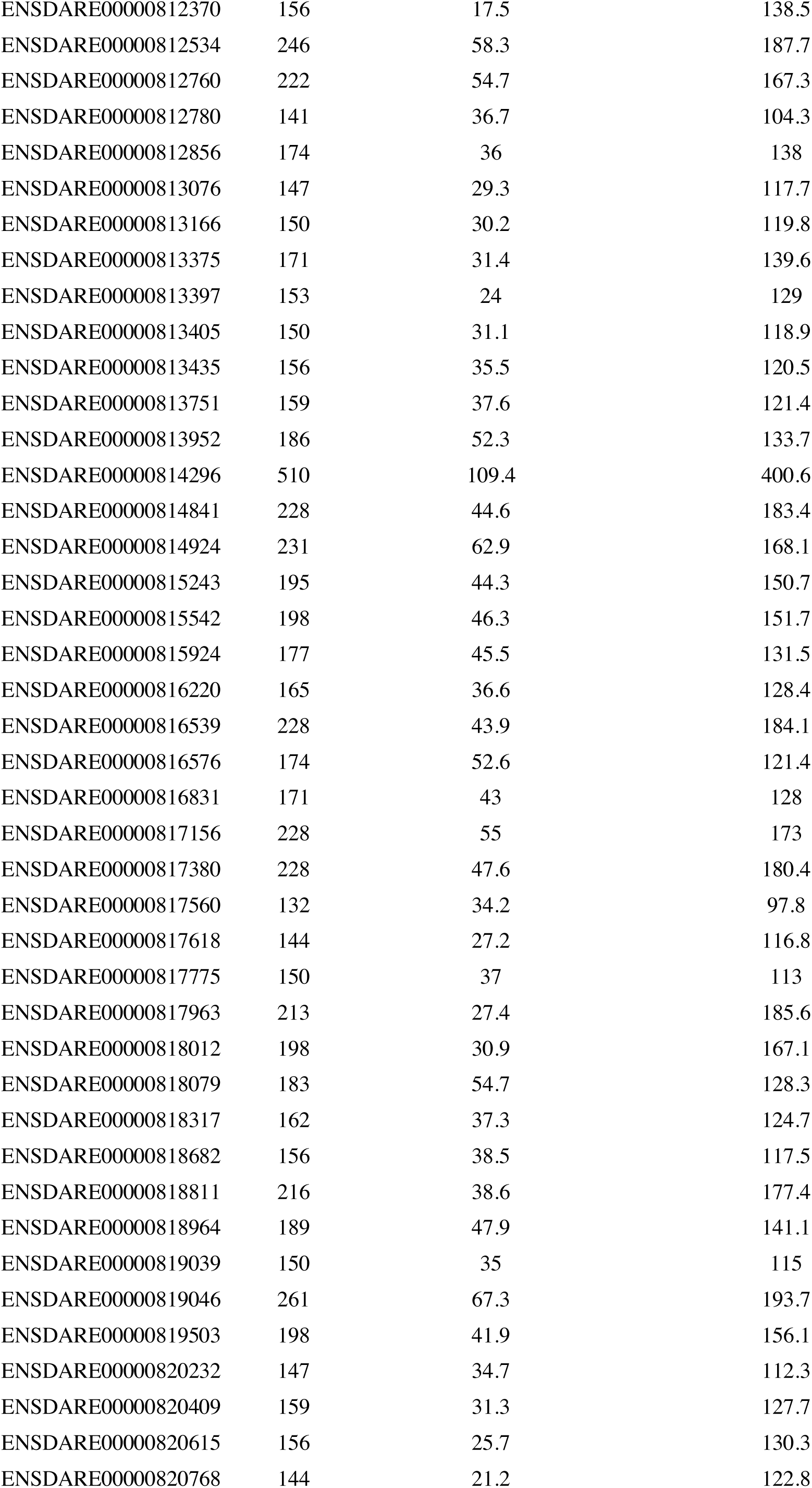
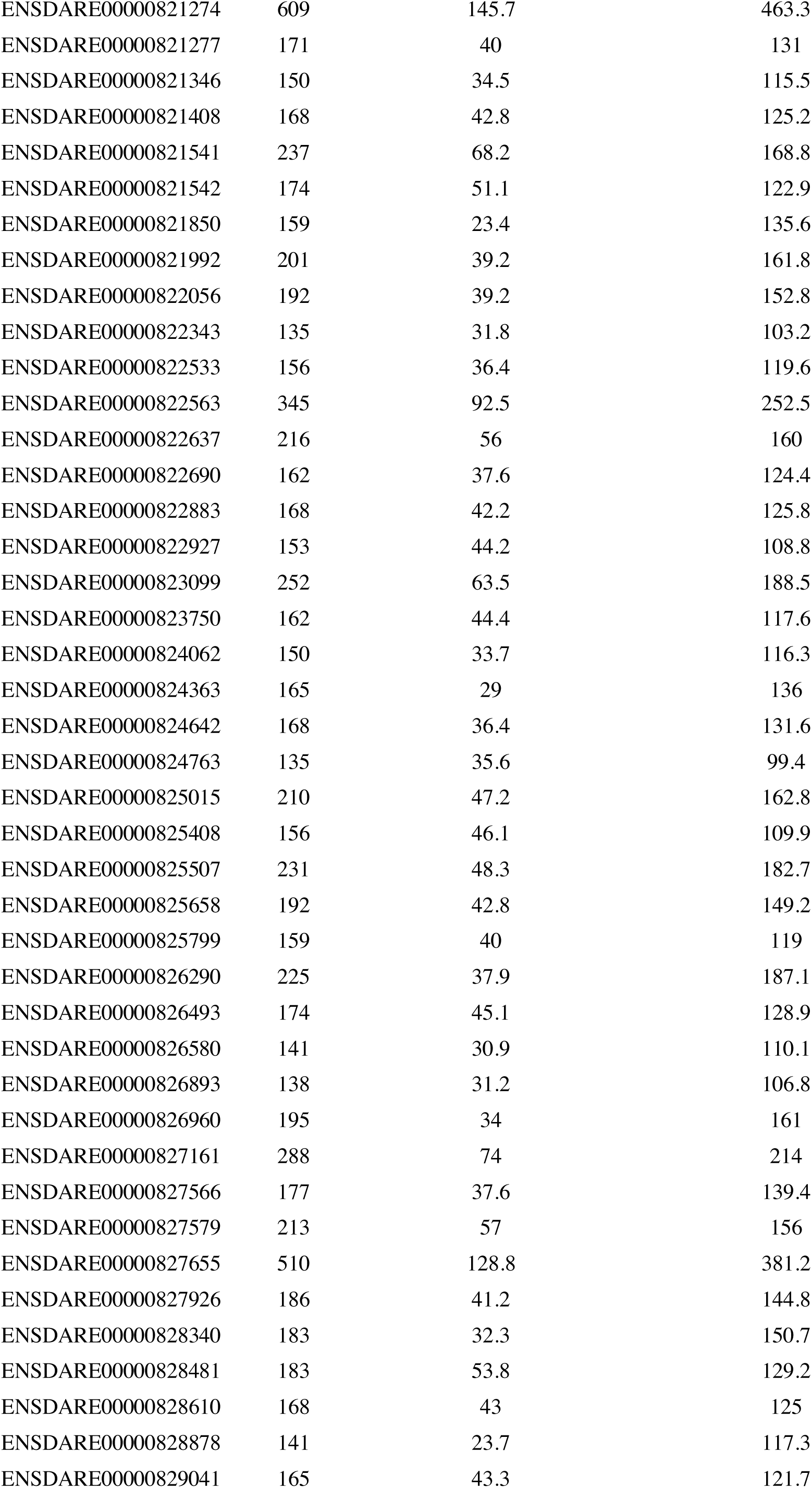
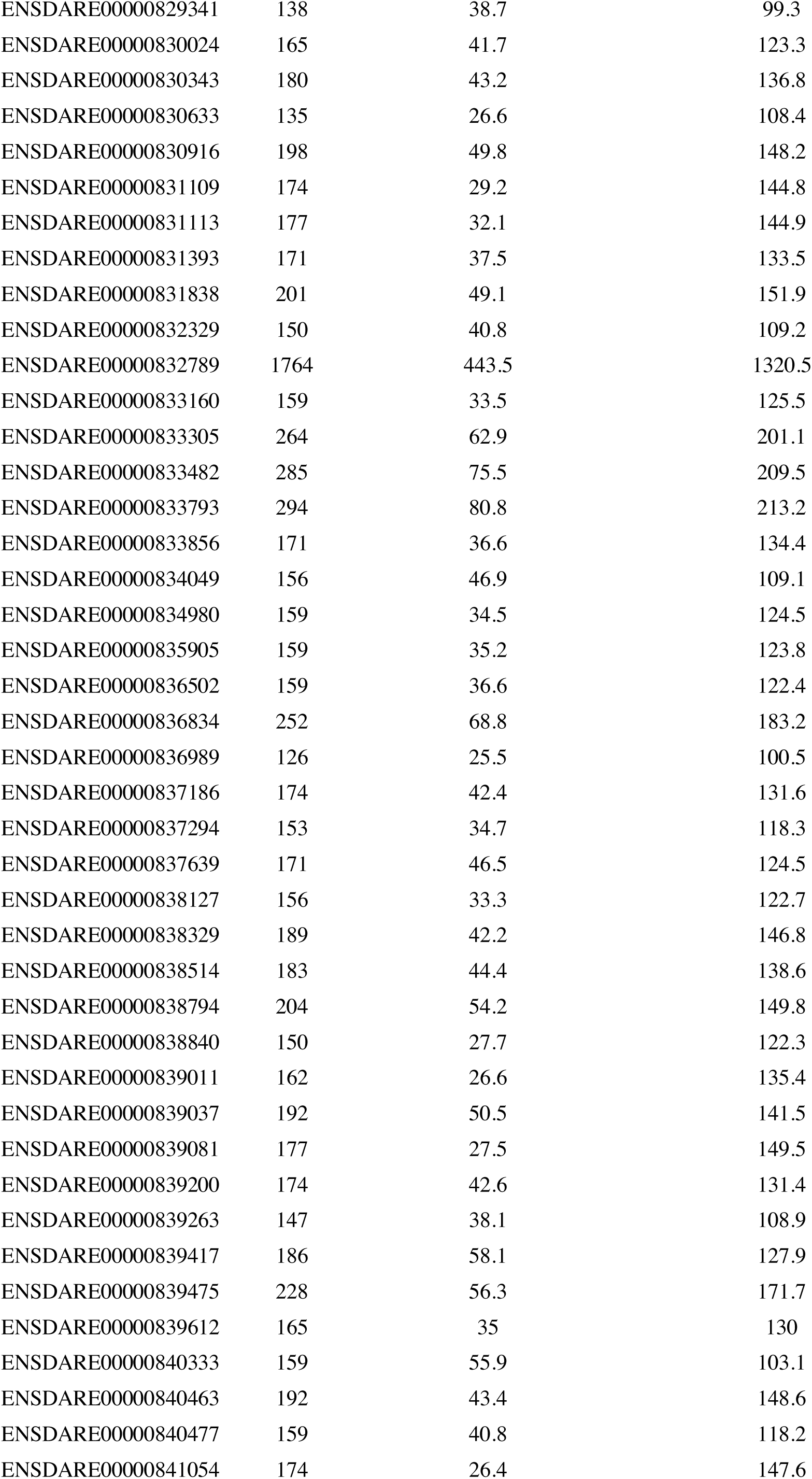
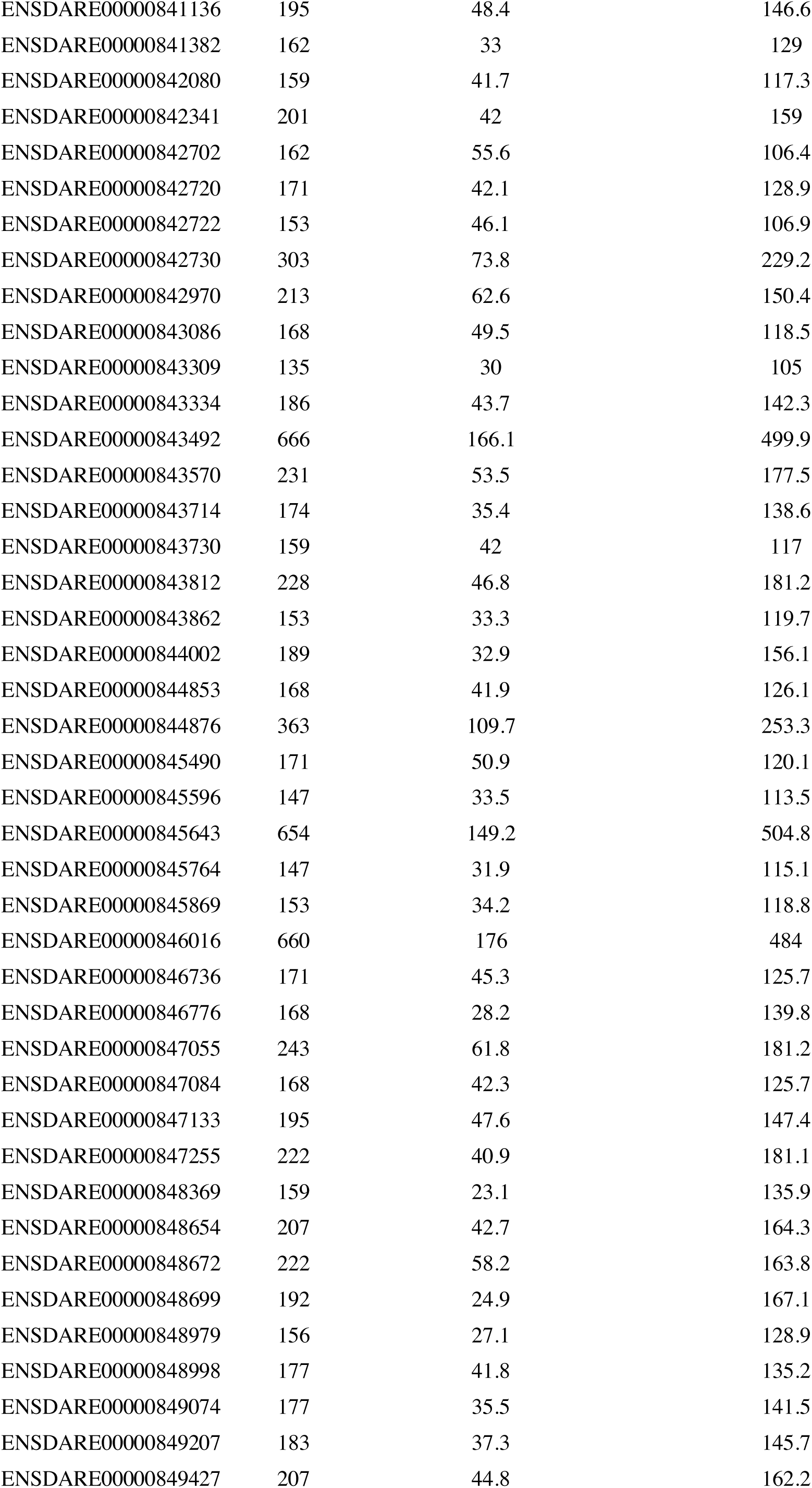
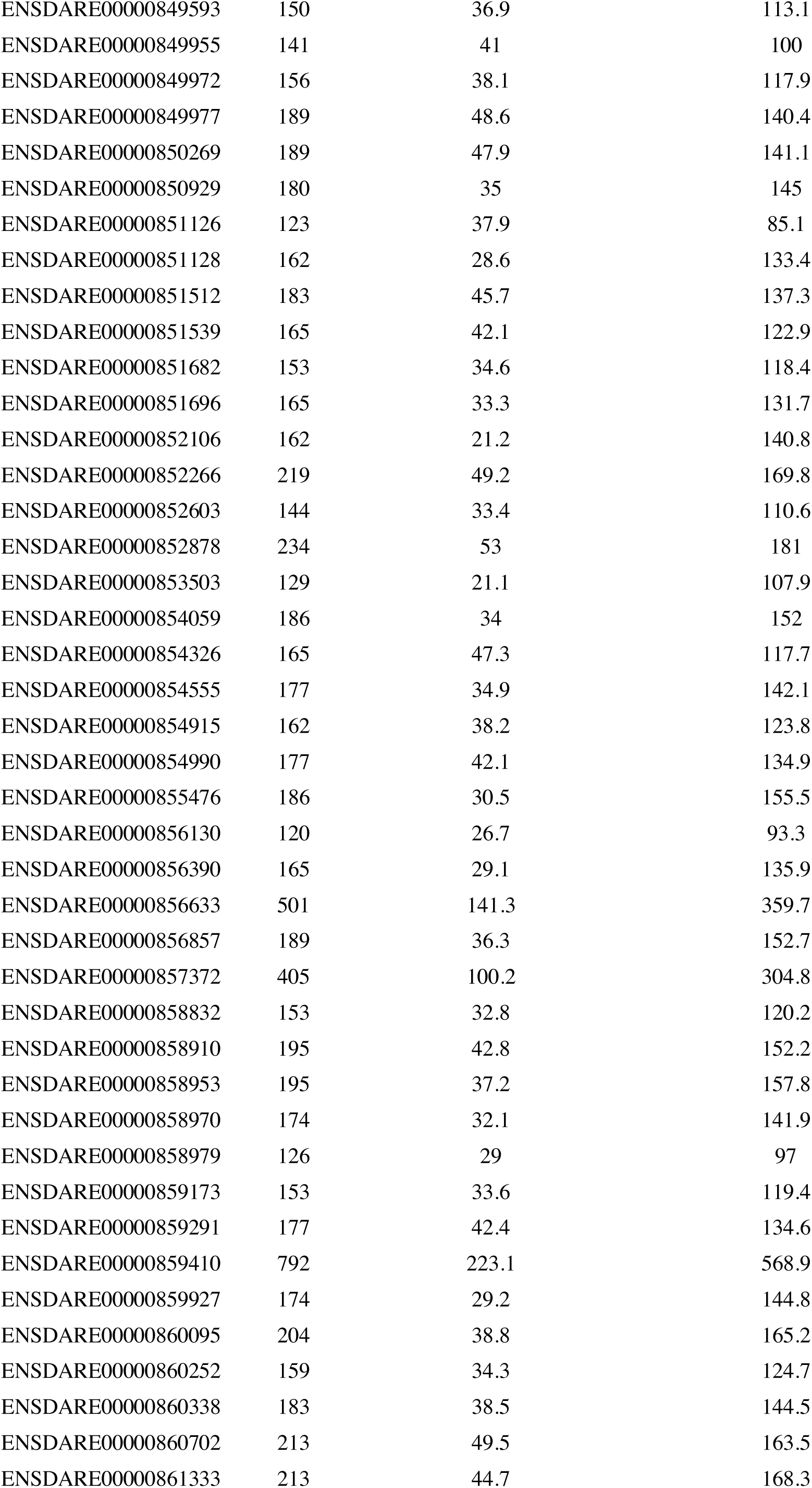
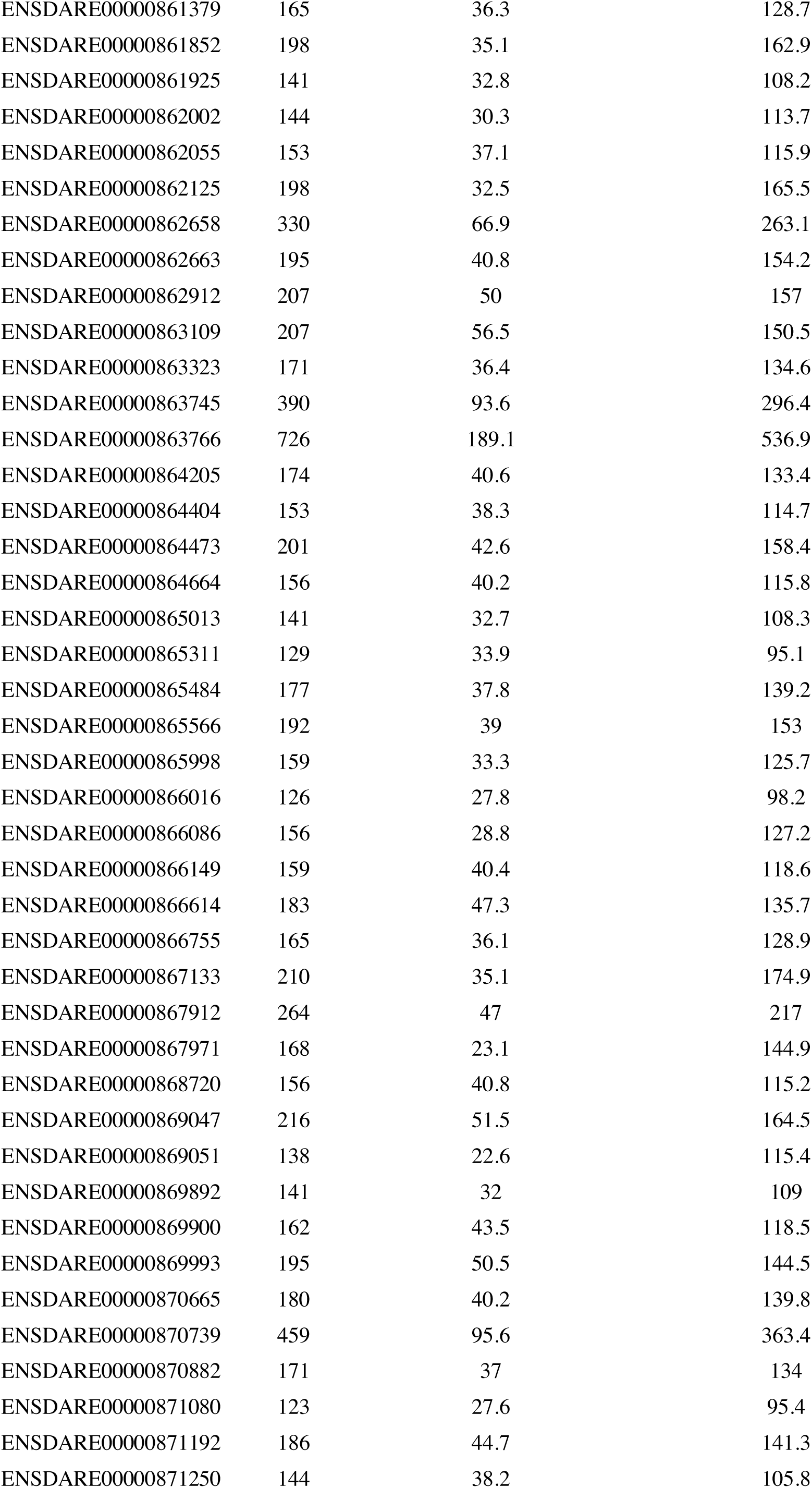
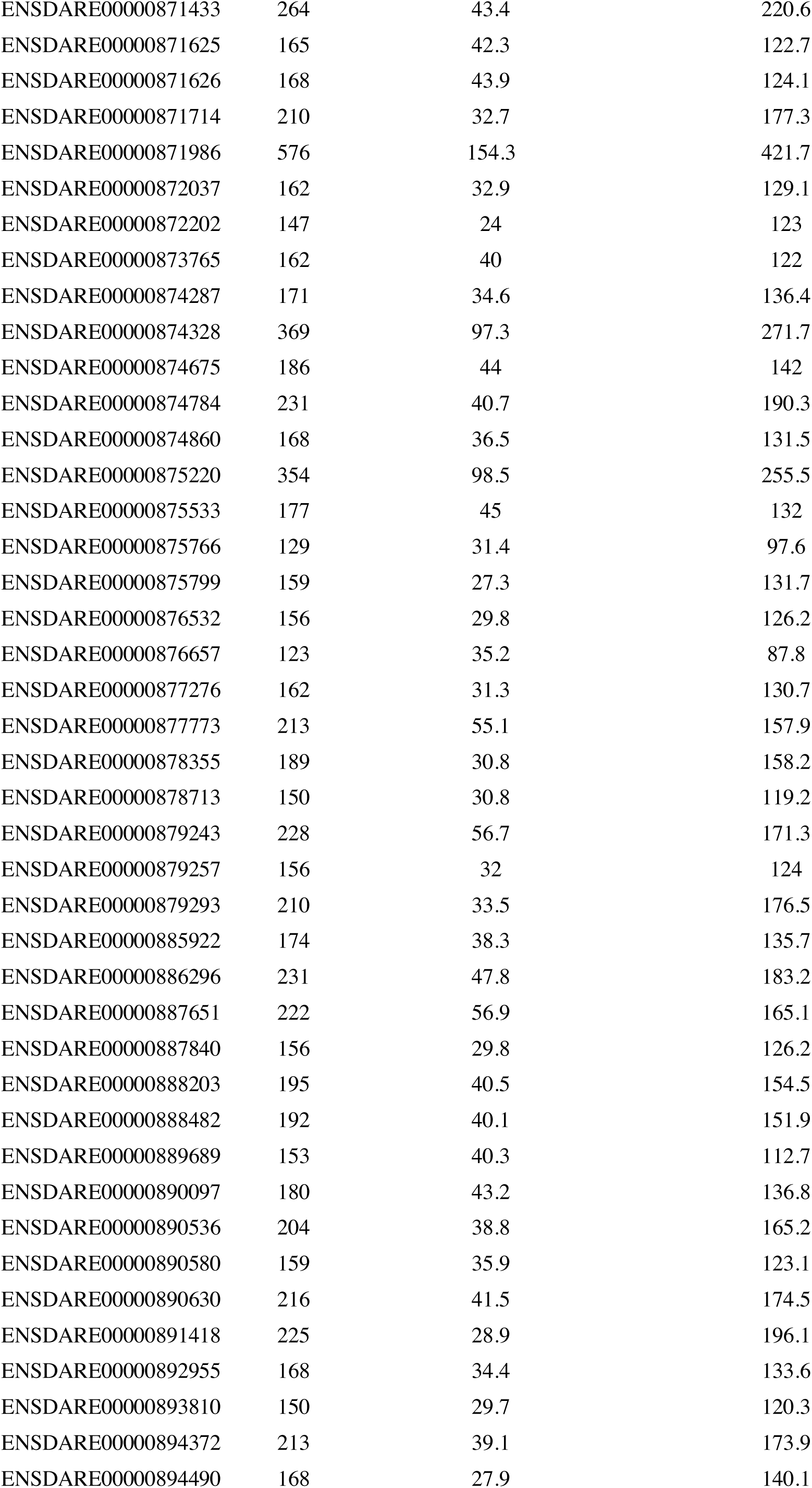
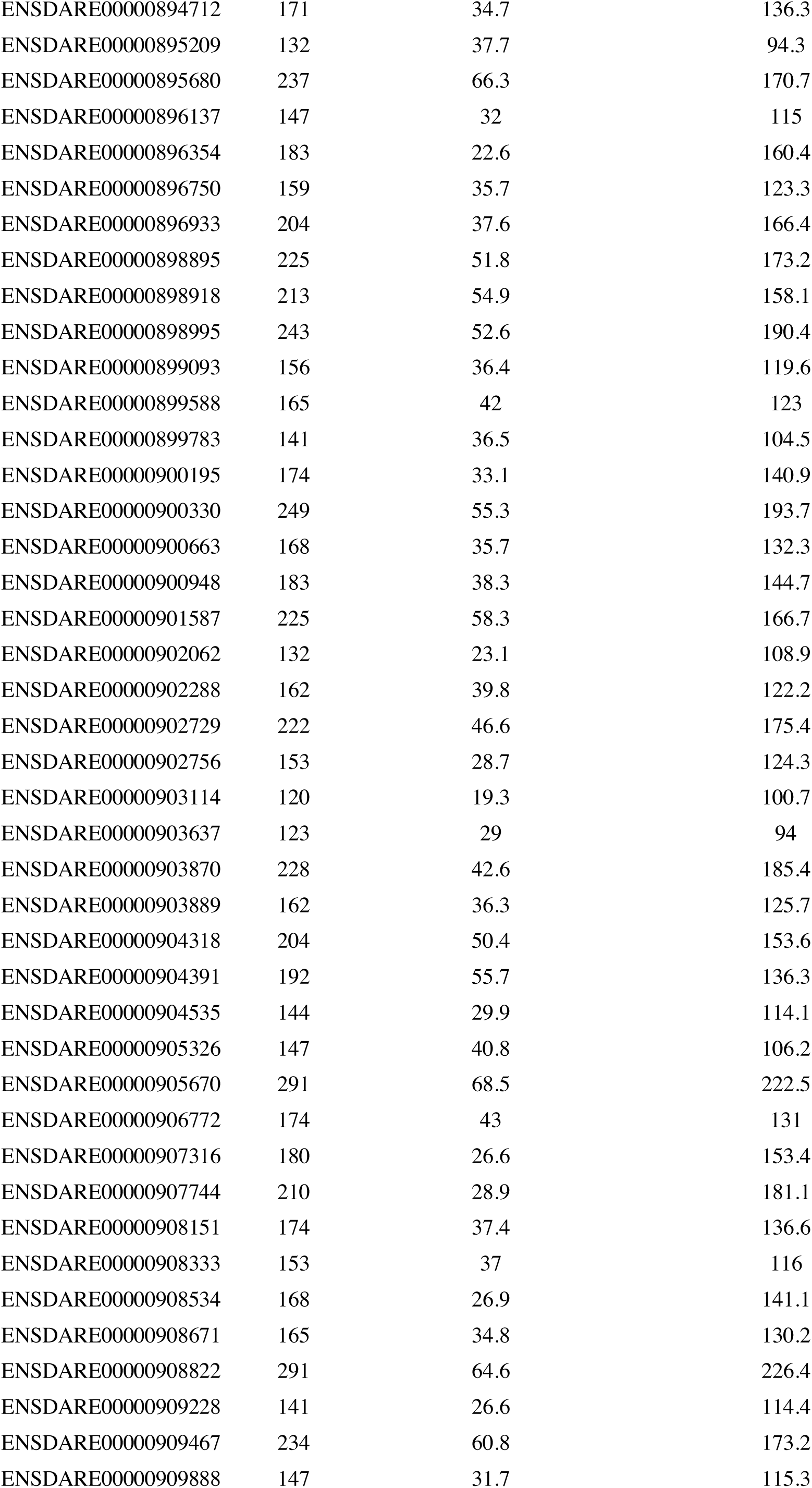
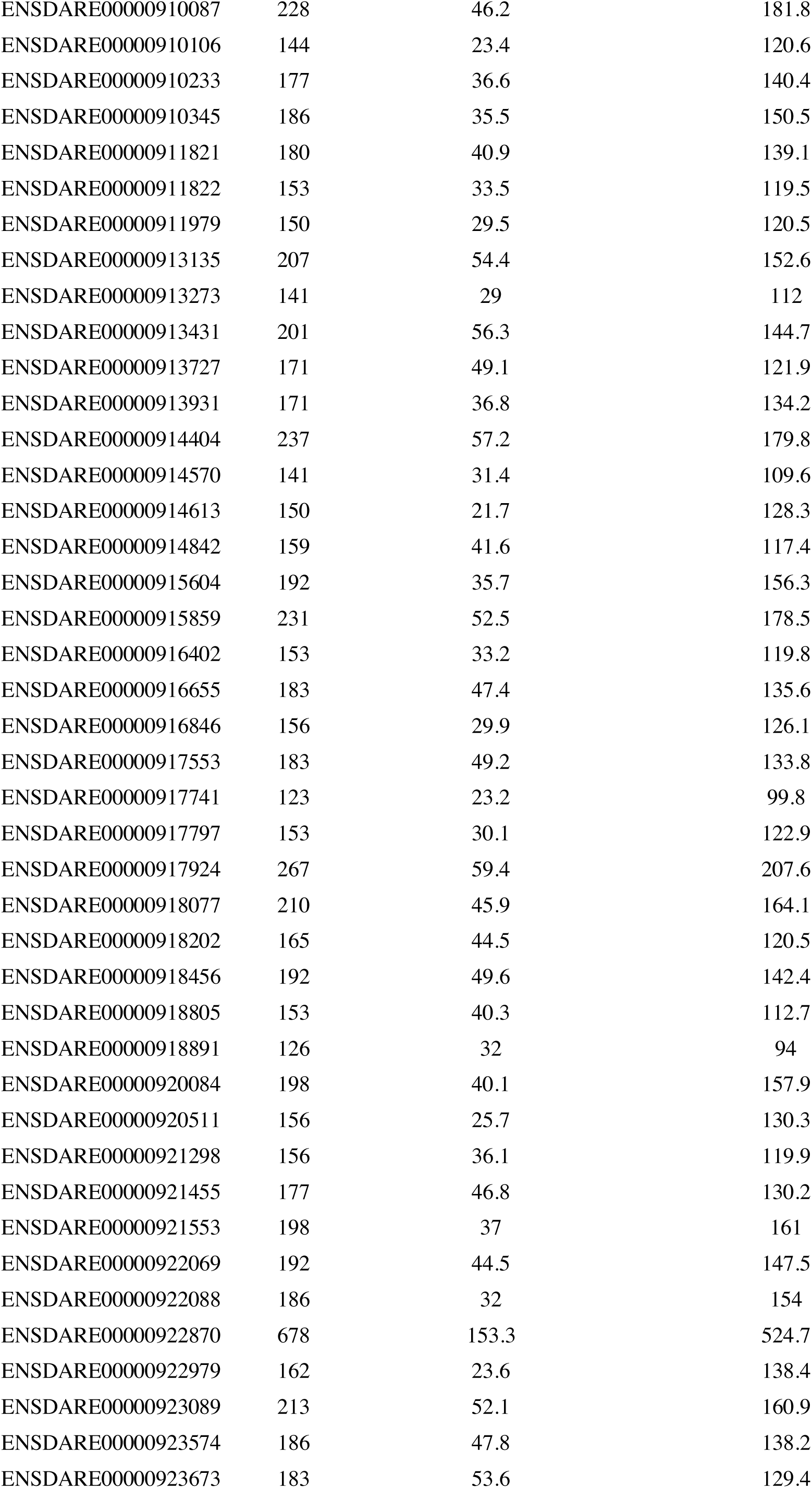
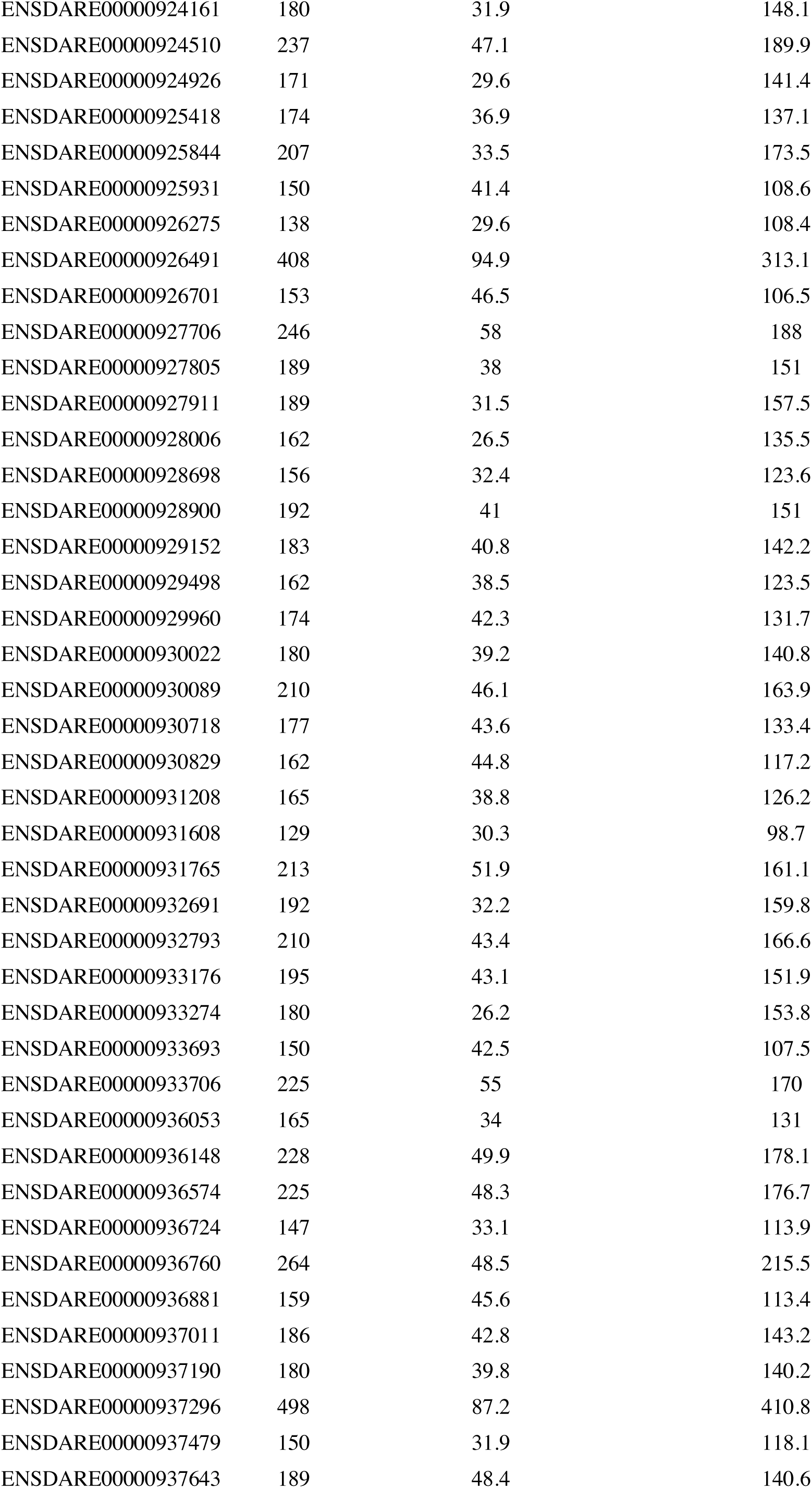
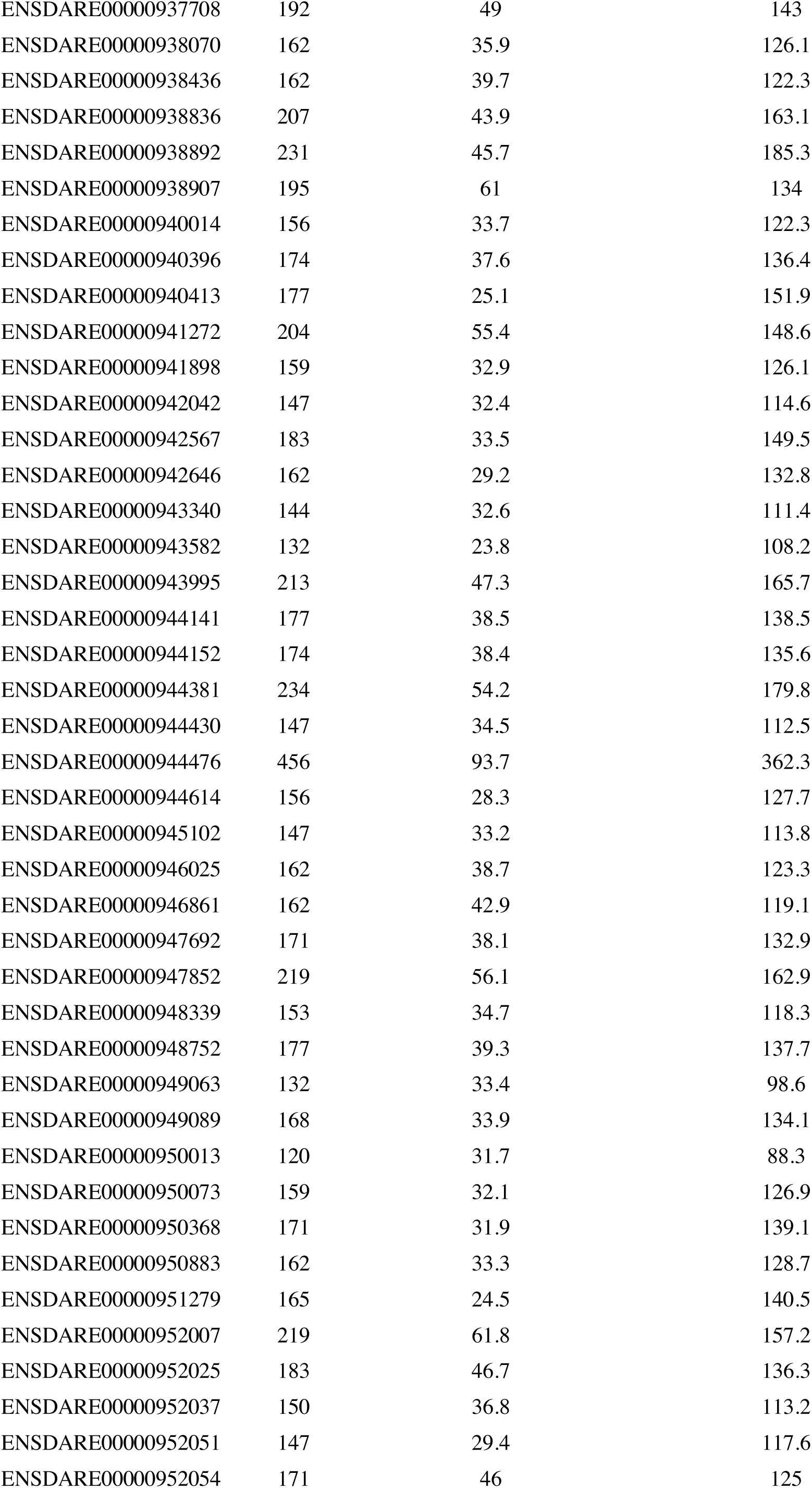
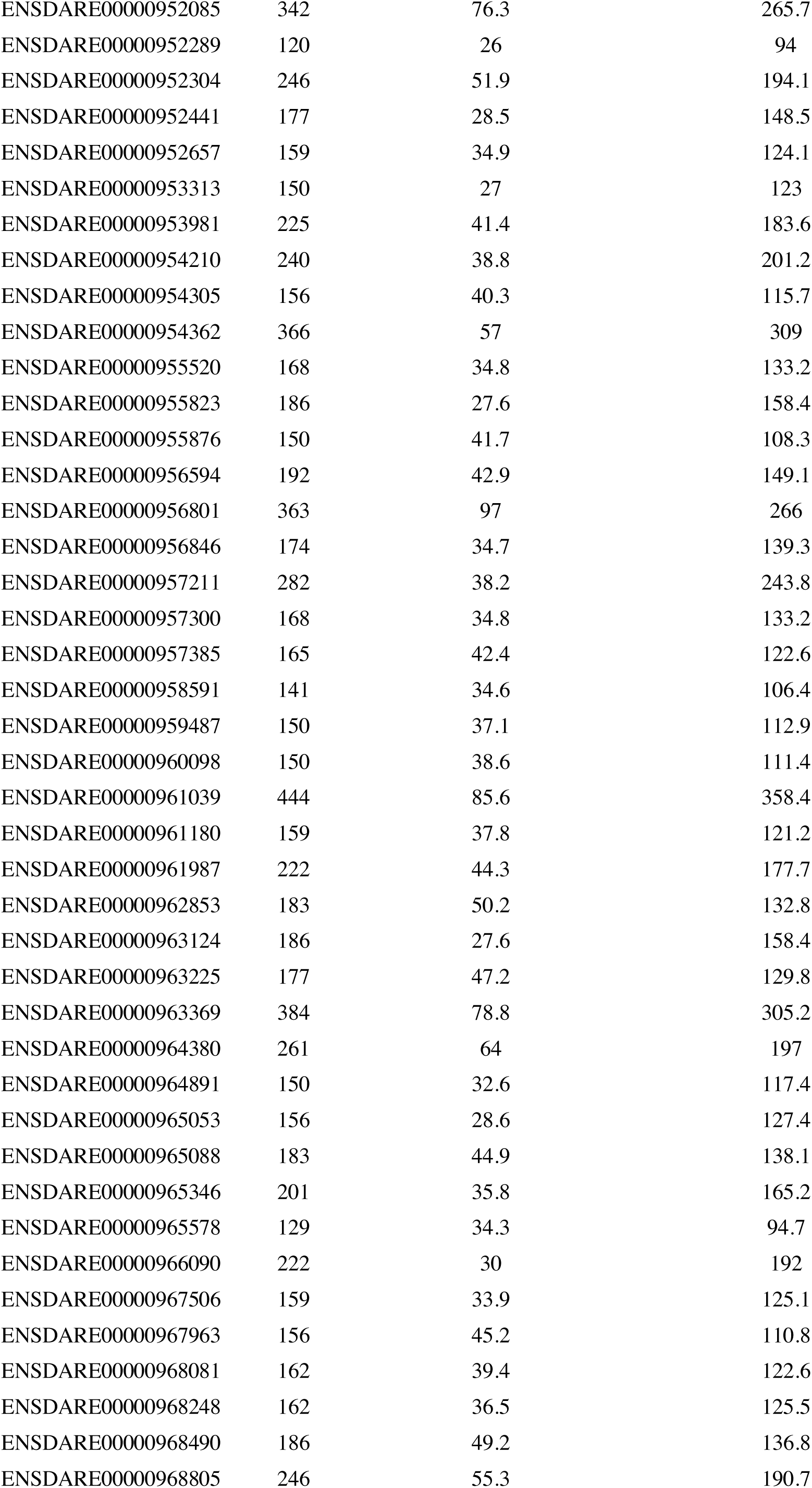
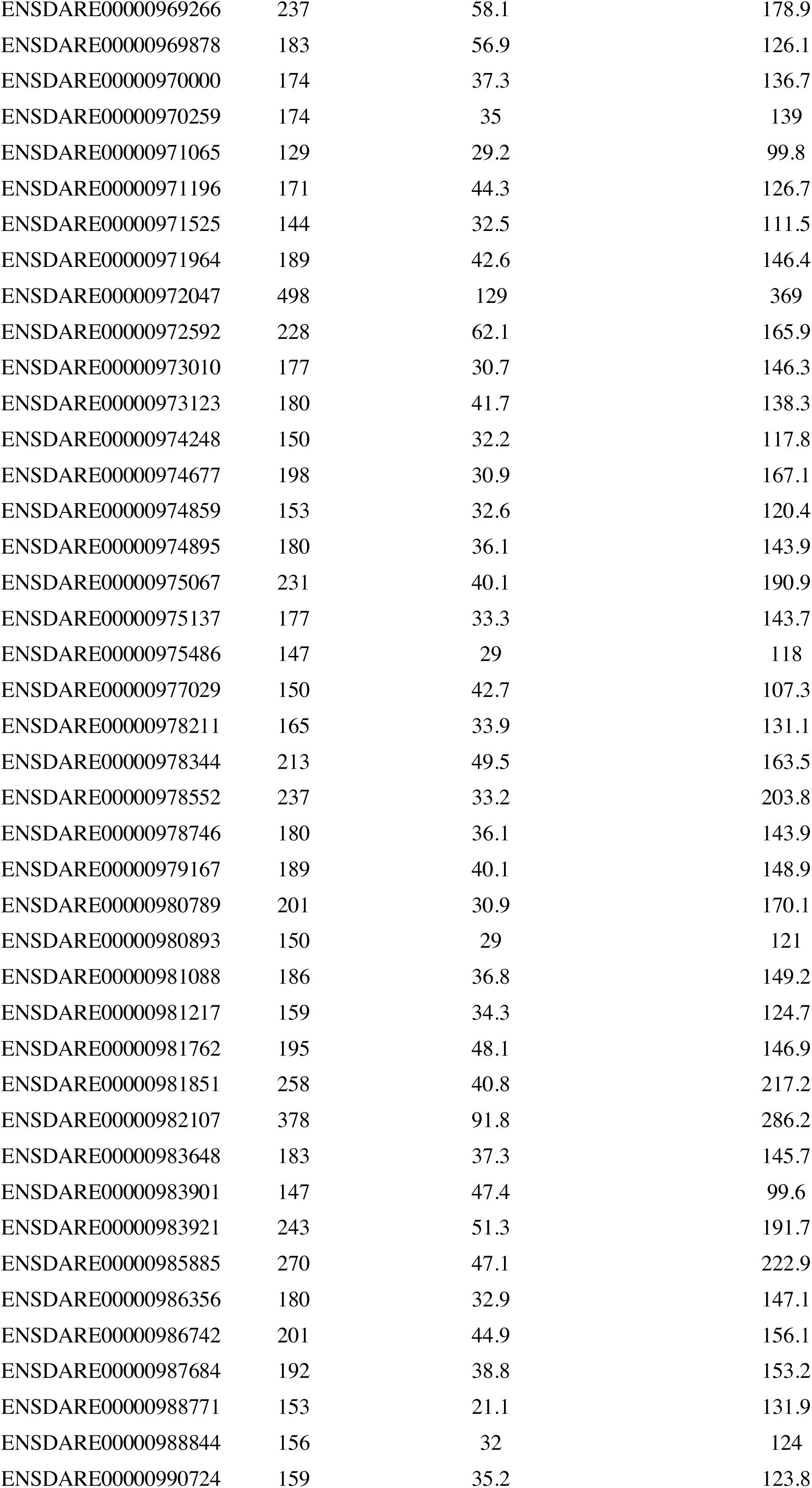
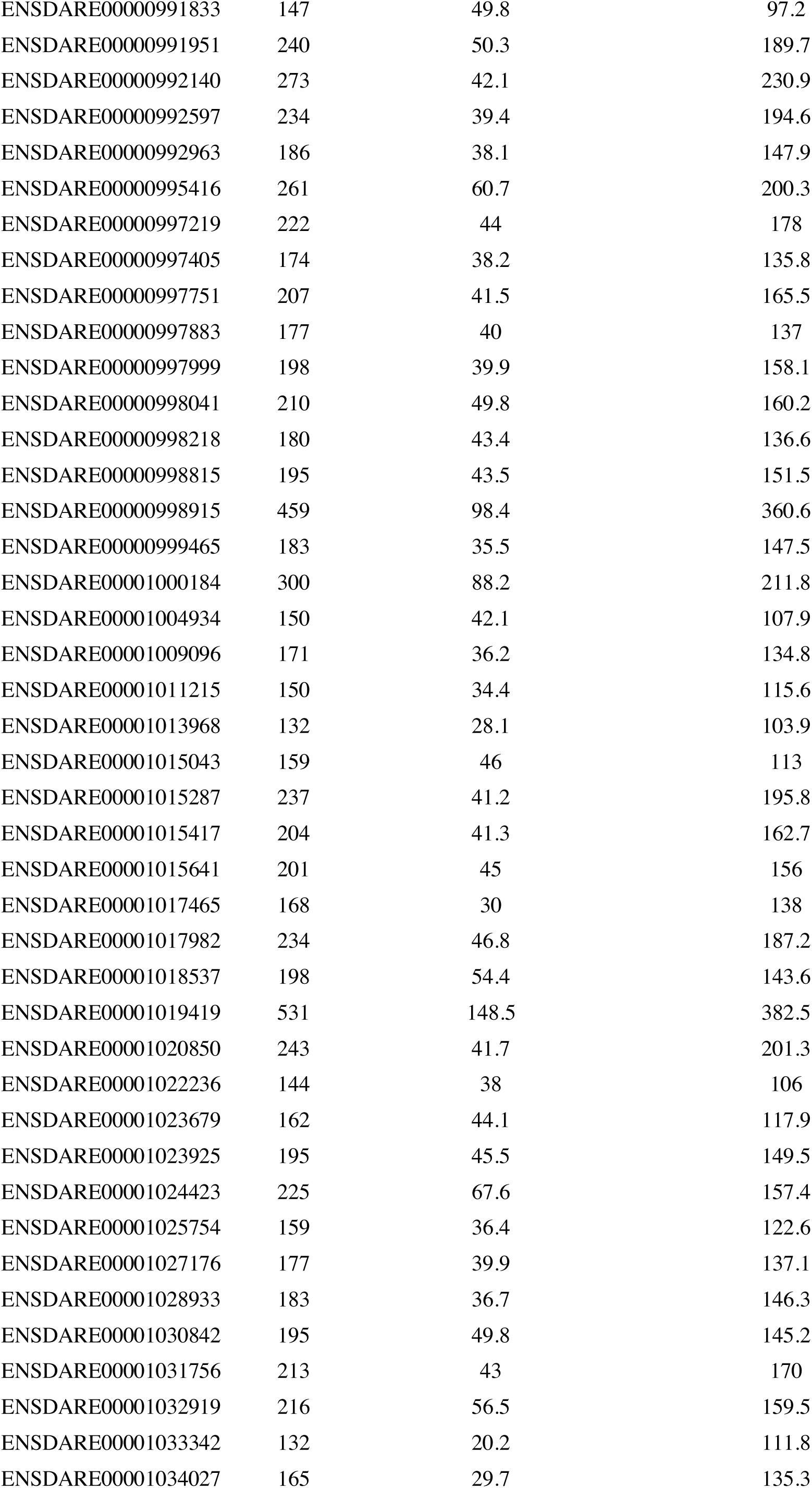
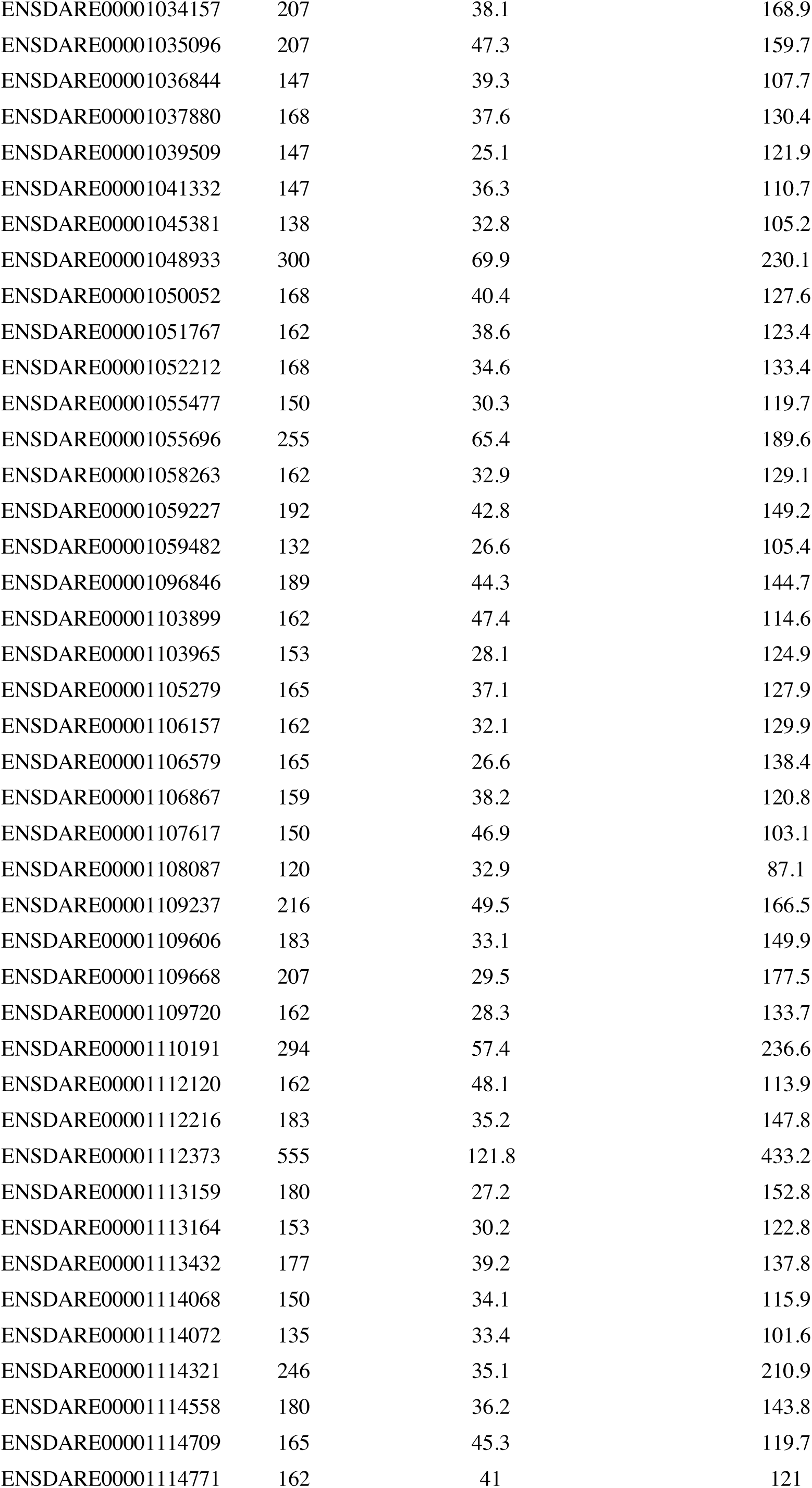
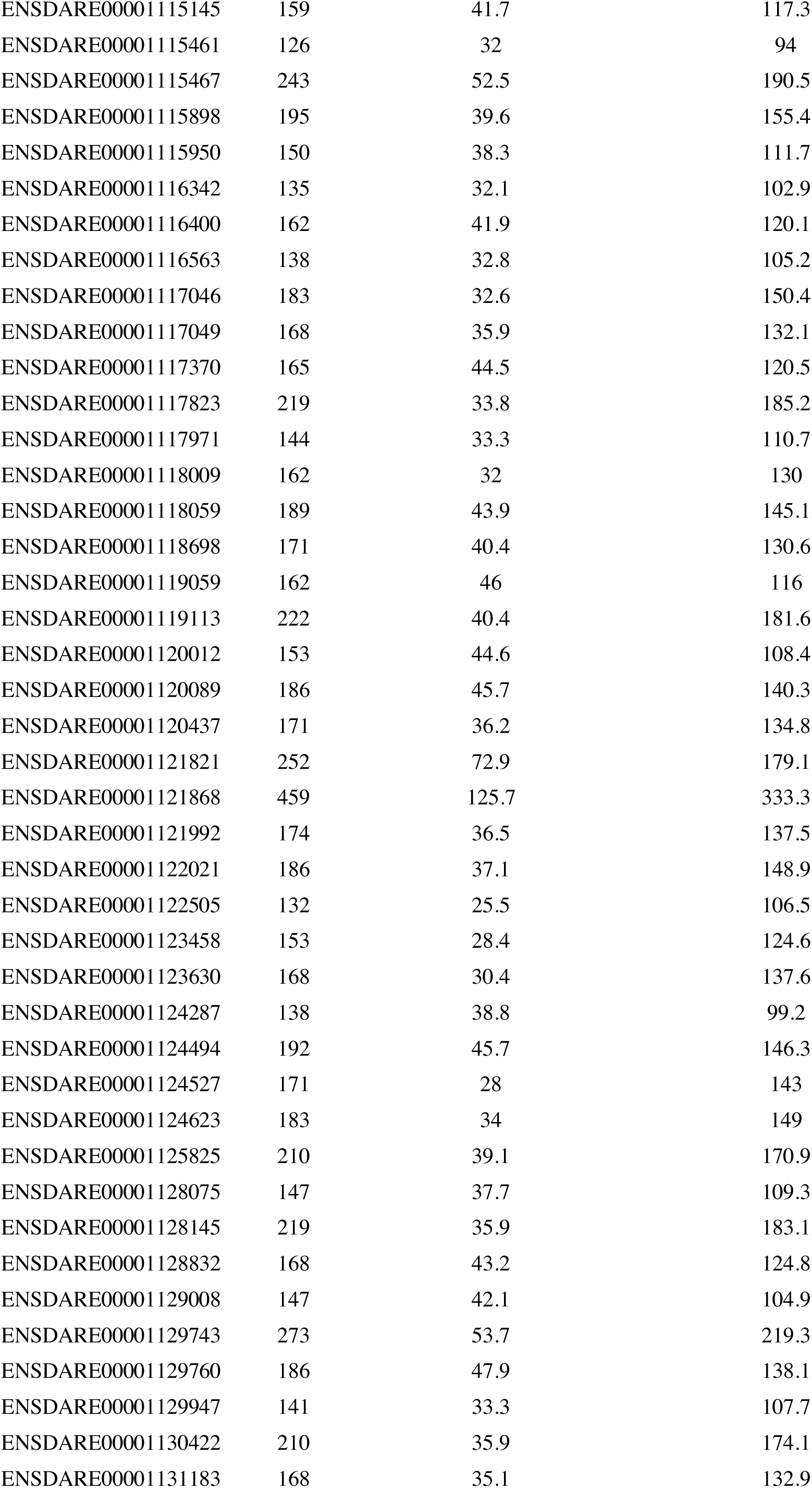
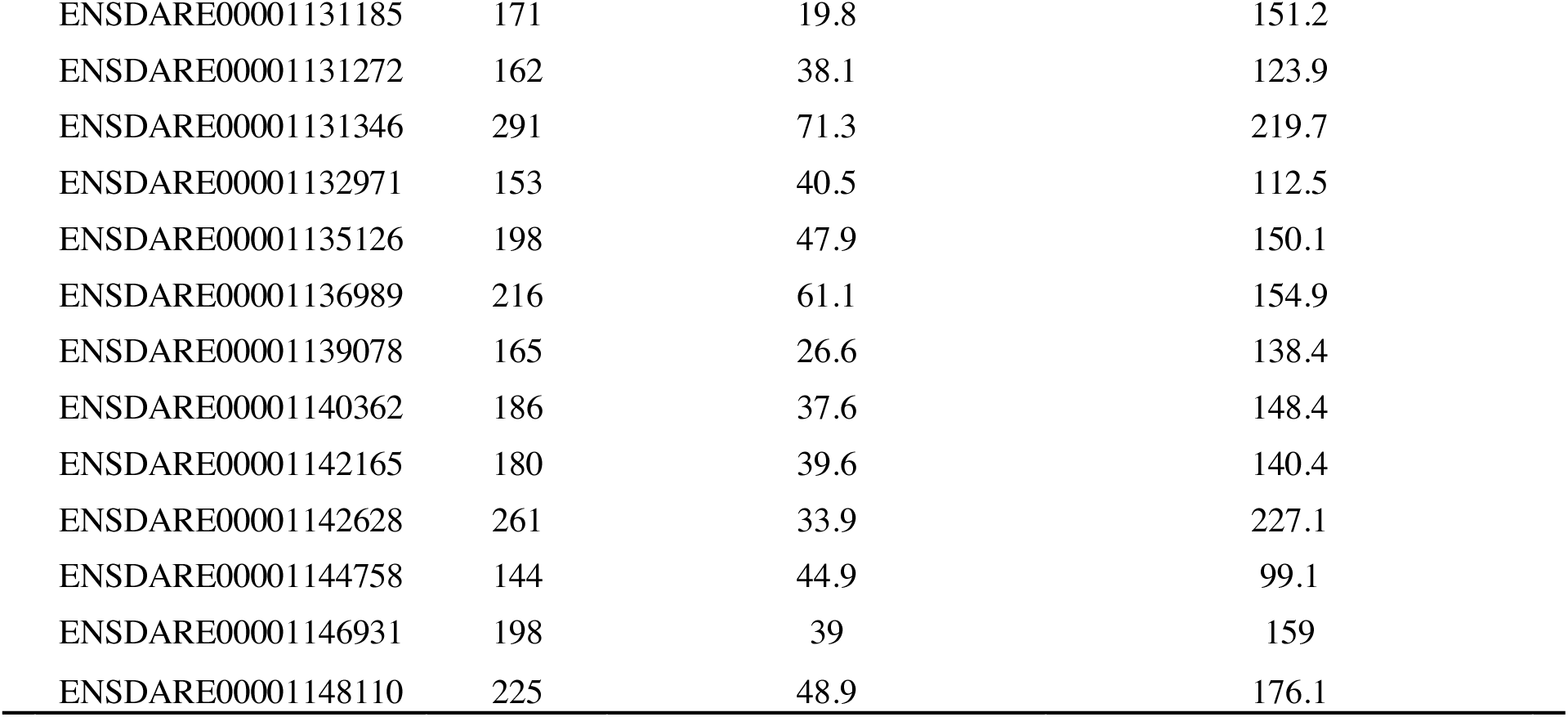
Information on the fish genomes. This table contains the detailed gene length in bp for each gene, the number of synonymous sites and the number of non-synonymous sites (number of synonymous sites + number of synonymous sites = gene length).

